# Characterization of MEK1/2 Degraders Uncovers a Kinase-Independent Role for MEK1/2 in the Stabilization and Maturation of CRAF

**DOI:** 10.1101/2025.03.11.642495

**Authors:** Jason S. Wasserman, Alison M. Kurimchak, Carlos Herrera-Montávez, Glenn A. Doyle, Brandon D. Fox, Ishadi K. M. Kodikara, Xiaoping Hu, Jianping Hu, Jian Jin, James S. Duncan

**Author notes:** These authors contributed equally (Jason S. Wasserman, Alison M. Kurimchak).

## Abstract

Altered MAPK signaling frequently occurs in human disease. MEK1 and MEK2 (MEK1/2) are central protein kinases in the MAPK signaling cascade that phosphorylate ERK1/2 promoting cell growth. MEK1/2 degraders offer a strategy to characterize both kinase-dependent and independent functions of MEK1/2. Here, we discovered that MEK1/2 degradation, but not kinase inhibition, caused the subsequent degradation of upstream kinase CRAF via a cell-intrinsic mechanism. MEK1/2 binding to CRAF, but not MEK1/2 catalytic activity, was required for CRAF protein stability and maturation to a functional kinase. In the absence of MEK1/2, a minor pool of newly synthesized immature CRAF that had anti-apoptotic functions remained. Finally, we showed that a stable primed CRAF-MEK1/2 signaling complex existed in cells that required RAS binding to potentiate MEK-ERK phosphorylation. Together, we’ve discovered a previously unrecognized kinase-independent function of MEK1/2, while contextualizing MEK1/2 as an integral component of the CRAF activation cycle beyond the conventional CRAF-MEK kinase-substrate paradigm.

## INTRODUCTION

The mitogen-activated protein kinase (MAPK) cascade is an evolutionarily conserved signaling pathway comprised of RAS GTPases, and downstream effector kinases (RAF, MEK, and ERK) that promote cell growth and survival (Bahar et al., 2023). The MAPK pathway is activated by growth factor receptor signaling inducing RAS GTPases to trigger activation of the RAF-MEK-ERK phosphorylation cascade, resulting in ERK-mediated regulation of various transcription factors that promote cell growth (Lavoie et al., 2020, Cargnello and Roux, 2012). Frequent alteration of the MAPK signaling cascade occurs in human disease, with ∼40% of all cancers exhibiting perturbed MAPK signaling (Santarpia et al., 2012). Moreover, activating mutations in key components of the MAPK cascade, such as KRAS or RAF kinases, are the underlying cause of several cancers. Consequently, the MAPK signaling pathway has been the focus of immense research leading to the development of inhibitors targeting the MAPK pathway. In particular, the MEK kinase has been a primary therapeutic target due to its central location within the MAPK pathway and highly druggable nature, with several MEK inhibitors currently being used to treat various MAPK-altered cancers.

MEK1 and MEK2 (MEK1/2) are ubiquitous members of the dual-specificity MAP2K kinase family that phosphorylate and activate ERK1/2 promoting cell growth and survival. MEK1/2 paralogs share highly similar kinase domains, including their activation loops, which are phosphorylated by the upstream RAF family of kinases ARAF, BRAF, and CRAF (Roskoski, 2012). MEK1/2 differ in their N-termini where MEK1 has a characterized ERK interaction. MEK1 uniquely contains an ERK negative feedback phosphorylation site within its proline-rich domain at T292 (Roskoski, 2012). Moreover, MEK1 but not MEK2 knockout is embryonic lethal, and MEK2 fails to compensate for MEK1 loss under specific settings (Bromberg-White et al., 2012), hinting at differential cellular roles of MEK1/2. Notably, although the canonical function of MEK1/2 in activating downstream kinase ERK1/2 is well documented, little is known about whether MEK1/2 exhibits other functions in cells.

The activation of MEK1/2 by RAF is thoroughly described; RAF directly interacts with and phosphorylates MEK1/2, resulting in MEK1/2 kinase activation and subsequent phosphorylation of ERK1/2. In contrast, how the upstream RAF kinase, particularly CRAF, is activated is highly complex, involving an orchestration of chaperones, phosphorylation, and RAF dimerization, ultimately resulting in the phosphorylation of MEK1/2. The recognized CRAF signaling cascade has MEK1/2 functioning solely as a substrate of CRAF, binding to CRAF after RAS binding and RAF dimerization. However, there is evidence that MEK1 can directly regulate CRAF activity independent of RAS (Leicht et al., 2013), suggesting that MEK1 has additional functions in the CRAF cycle. Here, using MEK1/2 proteolysis targeting chimeras (PROTACs), we discovered that MEK1/2 have kinase-independent functions in the regulation of CRAF protein stabilization, maturation, and kinase activation. Our findings reshape the current view of the CRAF activation cycle, establishing MEK1/2 binding to CRAF as an early required stabilizing event preceding CRAF-RAS engagement. Thus, MEK1/2 functions as both an activator and substrate of CRAF.

## RESULTS

### MEK1/2 Degradation but not Inhibition Caused Loss of Upstream Kinase CRAF

We performed global proteomics on PANC-1 cells treated with equal concentrations of the MEK1/2 inhibitor, PD0325901 or the PD0325901-based MEK1/2 degrader, MS934 (Hu et al., 2020). Strikingly, in addition to MEK1/2 protein loss following 24-hour MS934 treatment, we observed a significant reduction in total CRAF protein levels that was not detected in PD0325901- treated cells (**Fig. 1A-B, S1A, Data File S1**). Similarly, immunoblot analysis showed a dose-dependent reduction in total CRAF protein levels concomitant with MEK1/2 degradation in response to MS934 treatment of PANC-1 cells that was not observed with PD0325901 or a negative control PROTAC (MS934[-]) unable to recruit VHL (**Fig. 1C**). Quantitation of immunoblots revealed that CRAF total protein levels were reduced to ∼25% of the control by the 3.3 μM dose (**Fig. 1D**). Notably, ARAF and BRAF protein levels were not reduced by 24 hours MS934 treatment, with BRAF showing a modest increase in protein levels (**Fig. 1D**). No change in *RAF1, ARAF* or *BRAF* mRNA levels in PANC-1 cells was observed following 1 μM MS934 treatment, signifying that the reduction in CRAF protein level by MS934 occurred at the post-transcriptional level (**Fig. 1E**). Co-treatment of PANC-1 cells with MS934 and the proteasome inhibitor bortezomib restored MEK1, MEK2 and CRAF protein levels, demonstrating the involvement of the proteasome in the downregulation of CRAF by MS934 (**Fig. 1F**).

**Figure 1.**
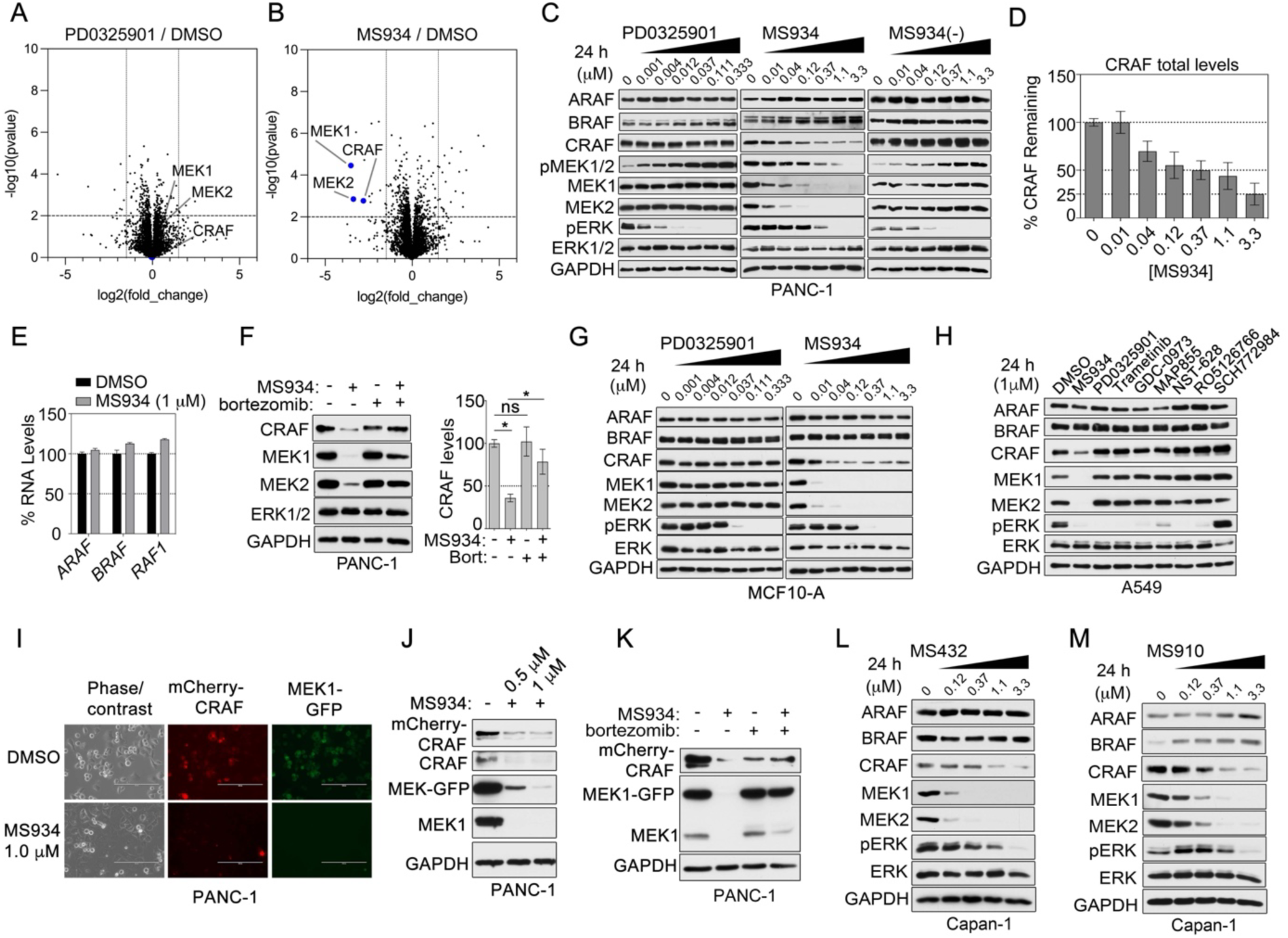
MEK1/2 degradation but not inhibition promotes CRAF loss. (A-B) Volcano plot of proteins increased or reduced in abundance in PANC-1 cells treated with PD0325901 (A) or MS934 (B) for 24 hours relative to DMSO-treated cells. Differences in protein log2 LFQ intensities among treated or control cells were determined by paired *t-*test permutation- based adjusted *P* values at FDR of <0.05 using Perseus software. (C-D) Immunoblotting for MAPK signaling components in PANC-1 cells treated with escalating doses of PD0325901, MS934, or MS934(-) for 24 hours. Blots are representative, and densitometric analysis of CRAF (D) are means ± SEM from three blots, each normalized to the loading control, GAPDH. (E) *ARAF*, *BRAF*, *RAF1* mRNA levels in PANC-1 cells following treatment with 1 μM MS934 for 24 hours, as determined by qRT-PCR. Data are means ± SEM of three independent experiments. (F) Immunoblotting for CRAF, MEK1/2, and ERK1/2 in PANC-1 cells treated with DMSO, 0.1 μM MS934, 0.1 μM bortezomib, or the combination for 24 hours. Cells were treated with bortezomib 2 hours prior to administering MS934. Blots are representative, and densitometric analysis of CRAF protein levels are means ± SEM from three blots, each normalized to the loading control, GAPDH. (G) Immunoblotting for MAPK signaling components in non-tumorigenic cell line MCF-10A treated with escalating doses of PD0325901, or MS934 for 24 hours. (H) Immunoblotting for MAPK signaling components in A549 cells treated with 1 μM MS934, MEK1/2 inhibitors, or ERK1/2 inhibitor for 24 hours. (I-J) Microscopy image (I) and immunoblot analysis (J) of mCherry-CRAF or MEK1-GFP expression in PANC-1 cells treated with 1 μM MS934 for 24 hours. Images are representative of three independent experiments. Scale bars, 200 μm. (K) Immunoblot analysis for mCherry-CRAF and MEK1-GFP in PANC-1 cells treated with DMSO, 1 μM MS934, 0.1 μM bortezomib, or the combination for 24 hours. Cells were treated with bortezomib 2 hours prior to administering MS934. (L-M) Immunoblot analysis of MAPK signaling components in Capan-1 cells treated with escalating doses of MS432 (H) or MS910 (I) for 24 hours.

To determine whether loss of CRAF by MS934 treatment occurred in other cell lines, we treated a panel of cancer cell lines harboring KRAS mutations (**Fig. S1B**), NF1-loss (**Fig. S1C**), BRAF mutation (**Fig. S1D**), as well as non-tumorigenic cells (**Fig. 1G, S1E**) with escalating doses of MS934. A dose-dependent reduction in CRAF total protein levels but not ARAF, or BRAF was observed with MS934 treatment but not PD0352901 in the non-tumorigenic cell line MCF10-A (**Fig. 1G**), and MS934 reduced CRAF levels across all cell lines tested (**Fig. S1B-E)**. Additionally, treatment of cells with other MEK1/2 inhibitors or an ERK1/2 inhibitor failed to reduce CRAF like MS934, demonstrating that loss of CRAF was specific to MEK1/2 degradation (**Fig. 1H**). Like PANC-1 cells, CRAF loss following MS934 treatment could be restored via blocking the proteasome in A549, and no change in *RAF1* mRNA levels was observed with escalating doses of MS934 in HCT-116 cells (**Fig. S1F-G**).

Next, we exogenously expressed mCherry-CRAF or Myc-CRAF in PANC-1 cells co-expressing MEK1-GFP, treated with MS934 for 24 hours, and measured CRAF protein levels by fluorescence imaging or immunoblotting. Notably, MS934 treatment reduced exogenous mCherry-CRAF and MEK1-GFP in PANC-1 cells (**Fig. 1I-J**). Similarly, MS934 treatment reduced exogenous Myc-CRAF levels in a dose-dependent manner in PANC-1 co-expressing MEK1-GFP (**Fig. S1H**). Blocking the proteasome using bortezomib in combination with MS934 restored mCherry-CRAF levels (**Fig. 1K**).

To determine whether other MEK1/2-targeting PROTACs caused downregulation of CRAF similar to MS934, we treated Capan-1 cells with the VHL-based PD0325901-PROTAC MS432 (Wei et al., 2019), or MS910 (Hu et al., 2020), which uses a CRBN-recruiting moiety. Notably, treatment of Capan-1 cells with escalating doses of MS432 (**Fig. 1L**), or MS910 (**Fig. 1M**) reduced MEK1/2 and CRAF protein levels, demonstrating that using either VHL or CRBN-recruiting moieties decreased both CRAF and MEK1/2 protein levels in cells. Thus, these data suggest MEK1/2 have kinase-independent functions, intrinsic to all cell types, beyond their described kinase-dependent function in activating the MAPK signaling cascade through ERK phosphorylation.

### MEK1/2 Are Required for Preserving CRAF Protein Stability

To explore the timing of CRAF protein loss in relation to MEK1/2 degradation by MS934, we treated PANC-1, NCI-H23, HCT-116, and A549 cell lines with 1 μM MS934 for 2, 4, 8, 12, 18, 24, 48, or 72 hours and performed immunoblotting. Degradation of MEK1 and MEK2 proteins occurred within the first 2 hours of MS934 treatment, however, CRAF protein levels were not reduced by 50% until 8 hours (**Fig. 2A-B**). Similarly, global proteomics showed that MEK1/2 but not CRAF protein were depleted following 4-hour MS934 treatment (**Fig. S2A, Data File S1**), requiring 24 hours of MS934 treatment to observe CRAF loss (**Fig. S2B**). Notably, minimal changes in ARAF or BRAF protein levels were observed across the 72-hour time course (**Fig. 2A**).

**Figure 2.**
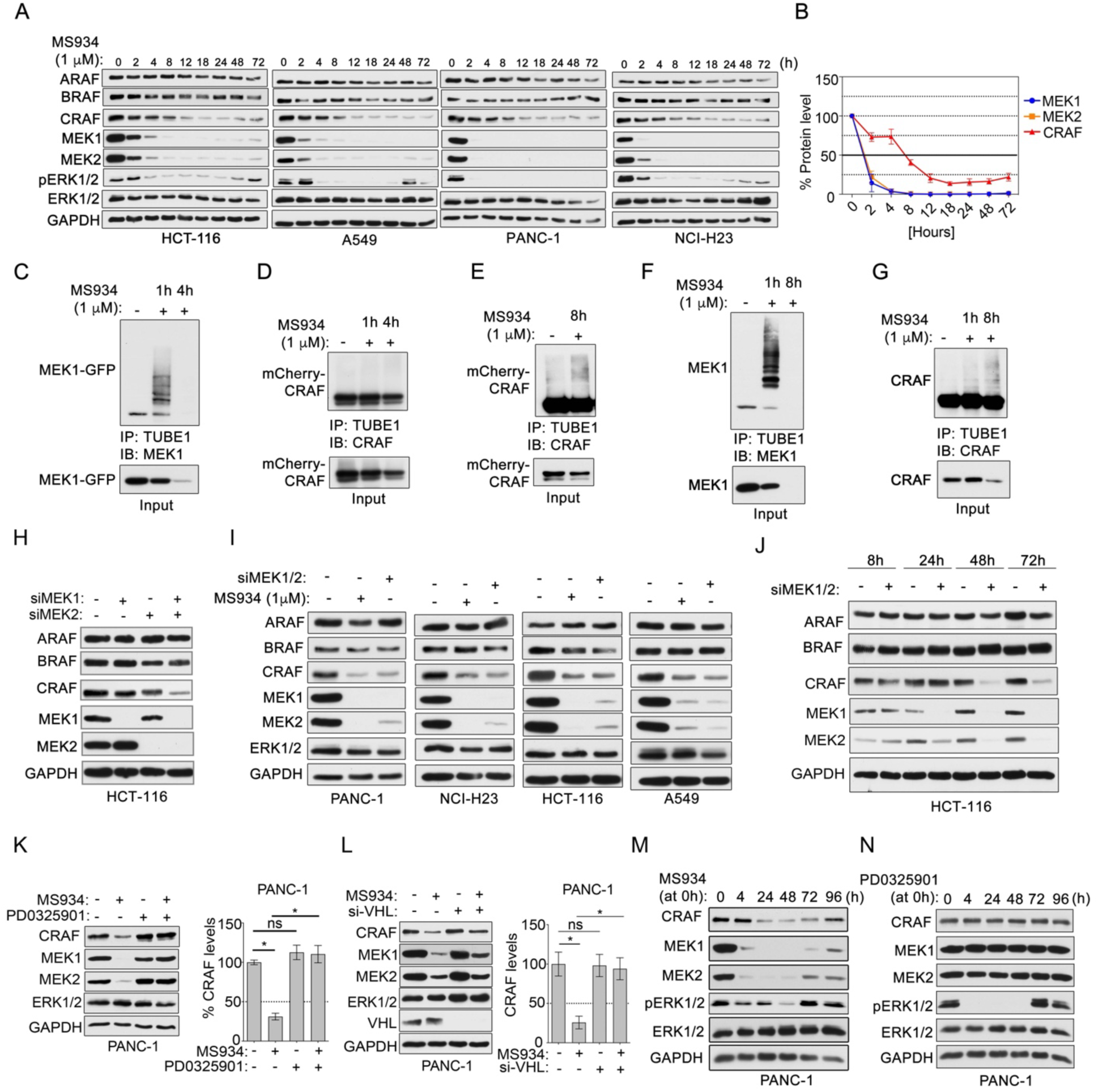
CRAF protein stability dependent on MEK1/2 expression in cells. (A-B) Immunoblot analysis of MAPK signaling in HCT-116, A549, PANC-1, and NCI-H23 following 1 μM MS934 treatment across a 72-hour time course. Fresh MS934 was administered every 24 hours. Blots are representative, and densitometric analysis of MEK1, MEK2, and CRAF (B) are means ± SEM across cell line blots, each normalized to the loading control, GAPDH. (C-G) Immunoprecipitation (IP) of ubiquitin-modified proteins using TUBE1 beads from PANC-1 cells co-expressing mCherry-CRAF and MEK1-GFP (C-E) or HCT-116 cells (F-G) treated with 1 μM MS934 for 1, 4 or 8 hours. The TUBE1 IP elutes were blotted with indicated antibodies. (H) Immunoblot analysis of RAF and MEK1/2 protein levels in HCT-116 cells treated with control siRNAs or MEK1, MEK2, or MEK1 and MEK2 siRNAs and cultured for 72 hours. (I) Immunoblot analysis of RAF and MEK1/2 levels in PANC-1, NCI-H23, HCT-116 and A549 treated with 1 μM MS934 or MEK1 and MEK2 siRNAs (siMEK1/2). (J) Immunoblot analysis of RAF and MEK1/2 levels in HCT-116 cells treated with siRNAs targeting both MEK1 and MEK2 for 8, 24, 48, and 72 hours. (K) Immunoblotting for CRAF, MEK1/2, or ERK1/2 in PANC-1 cells treated with DMSO, 1 μM PD0325901, 0.1 μM MS934, or the combination for 24 hours. Cells were treated with PD0325901 2 hours prior to adding MS934. Blots are representative, and densitometric analysis of CRAF levels are means ± SEM from three blots, each normalized to the loading control, GAPDH. (L) Immunoblot analysis for CRAF, MEK1/2 or ERK1/2 in PANC-1 cells transfected with control siRNA or siRNAs targeting VHL and treated with DMSO or 0.1 μM MS934 for 24 hours. Blots are representative, and densitometric analysis of CRAF protein levels are means ± SEM from three blots, each normalized to the loading control, GAPDH. (M-N). Immunoblot analysis of MAPK signaling in PANC-1 cells treated with a single dose of 0.1 μM MS934 (M) or PD0325901 (N) and followed across a 96-hour period. No drug replenishment was performed.

To further characterize the timing and process of CRAF degradation following MS934 treatment, we measured ubiquitination of exogenous MEK1-GFP and mCherry-CRAF in PANC-1 cells in response to MS934 using TUBE immunoprecipitations (TUBE-IP), a method for enriching proteins containing poly-ubiquitination (Hjerpe et al., 2009). Analysis of MS934- mediated ubiquitination using TUBE1-IP, showed that MEK1-GFP was ubiquitinylated at 1 hour after 1 μM MS934 treatment, while CRAF showed no evidence of ubiquitination at 1 or 4 hours post-MS934 treatment (**Fig. 2C-D**). However, consistent with immunoblot analysis, TUBE1-IP at the later 8-hour time point showed ubiquitination of mCherry-CRAF following 1 μM MS934 treatment (**Fig. 2E**). Similarly, TUBE-IP of endogenous MEK1 or CRAF from HCT-116 cells following 1 μM MS934 treatment showed that ubiquitination of endogenous MEK1 but not CRAF was observed at 1-hour post-MS934 treatment, with CRAF showing ubiquitination at 8 hours post- MS934 treatment (**Fig. 2F-G**).

Next, we performed genetic knockdown of MEK1, MEK2 or both using siRNAs in HCT116 cells. Genetic depletion of MEK1 or MEK2 alone did not reduce CRAF protein levels, whereas the simultaneous knockdown of both MEK1/2 reduced CRAF levels, establishing that both MEK1 and MEK2 were required for maintaining CRAF protein levels (**Fig. 2H, S2C**). Whereas knockdown of MEK1/2 reduced CRAF, as observed with MS934 treatment, ARAF or BRAF protein levels were unaffected by MEK1/2 knockdown in PANC-1, NCI-H23, HCT-116 and A549 cells (**Fig. 2I**). MEK1/2 knockdown over a 72-hour period in HCT116 cells revealed that MEK1/2 proteins were reduced at 24 hours, while CRAF total protein levels remained until 48 hours, demonstrating MEK1/2 reduction preceded CRAF loss following MEK1/2 knockdown (**Fig. 2J**). Notably, knockdown of CRAF did not reduce MEK1/2 protein levels (**Fig. S2D**), and genetic depletion of the KSR1/2 scaffold proteins or ERK1/2 also failed to decrease CRAF protein levels as was observed with MEK1/2 knockdown (**Fig. S2E**). No changes in mRNA levels of *RAF1* were detected following MEK1/2 knockdown (**Fig. S2F**).

Next, we evaluated whether preventing MS934-mediated degradation of MEK1/2 would rescue CRAF protein levels using excess PD0325901 or performing knockdown of E3 ligase components VHL or CUL2 required by MS934. We observed restoration of MEK1/2 proteins in PANC-1 or NCI-H23 cells by combining MS934 with excess PD0325901 treatment (**Fig. 2K, S2G**). Similarly, preventing PROTAC-mediated degradation via VHL knockdown (**Fig. 2L, S2H**) blocked CRAF degradation, restoring CRAF protein levels. Additionally, knockdown of CUL2 in MS934-treated HCT-116 cells restored MEK1/2 and CRAF protein levels (**Fig. S2I**). Finally, analysis of CRAF and MEK1/2 protein levels in PANC-1 cells following a single treatment of MS934 over a 96-hour time course, showed that CRAF protein levels returned to control levels by 96 hours coinciding with the partial return of MEK1 and MEK2 (**Fig. 2M**). No changes in CRAF or MEK1/2 protein levels were detected over 96 hours following PD0325901 treatment (**Fig. 2N**). Taken together, these data demonstrate that degradation of MEK1/2 promoted the subsequent degradation of CRAF in KRAS mutant cells several hours after MS934 treatment or MEK1/2 genetic knockdown by a proteosome-dependent process, suggesting that CRAF protein stability was dependent on the presence of MEK1/2 proteins.

### MEK1/2 Degradation Caused Rapid Dephosphorylation of CRAF at S621 Promoting Misfolding and Degradation

CRAF activation is regulated by several phosphorylation events (Dhillon et al., 2007, Noble et al., 2008, Reiterer et al., 2010, Yu et al., 2023, Xiang et al., 2002, Zang et al., 2008). To explore the effect of MEK1/2 degradation on CRAF phosphorylation, we performed immunoblot analysis of HCT-116 cells treated with 1 μM MS934 across a 72-hour time course (**Fig. 3A-B**). Reduction of p-S289/296/301 was observed within 2 hours of MS934 treatment, consistent with rapid inhibition of ERK1/2 signaling (Dougherty et al., 2005). Whereas CRAF p-S338 levels varied in HCT-116 cells over time, with levels being reduced at later time points, decreased p-S338 levels were observed within 4 hours of MS934 treatment in A549, PANC-1, and NCI-H23 cell lines, signifying inhibition of CRAF activity (King et al., 1998). CRAF p-S259 was downregulated within 2 hours post-MS934 in all cell lines. Autophosphorylation of S621 by CRAF is essential for protein stability, where mutation of S621 to alanine promotes CRAF degradation (Noble et al., 2008). Notably, decreased levels of p-S621 were observed 2 hours post-MS934 treatment in all cells, preceding CRAF protein degradation. Reduced CRAF p-S621 was also observed in KRAS wild-type HEK293T cells in response to 4-hour MS934 treatment (**Fig. S3A**).

**Figure 3.**
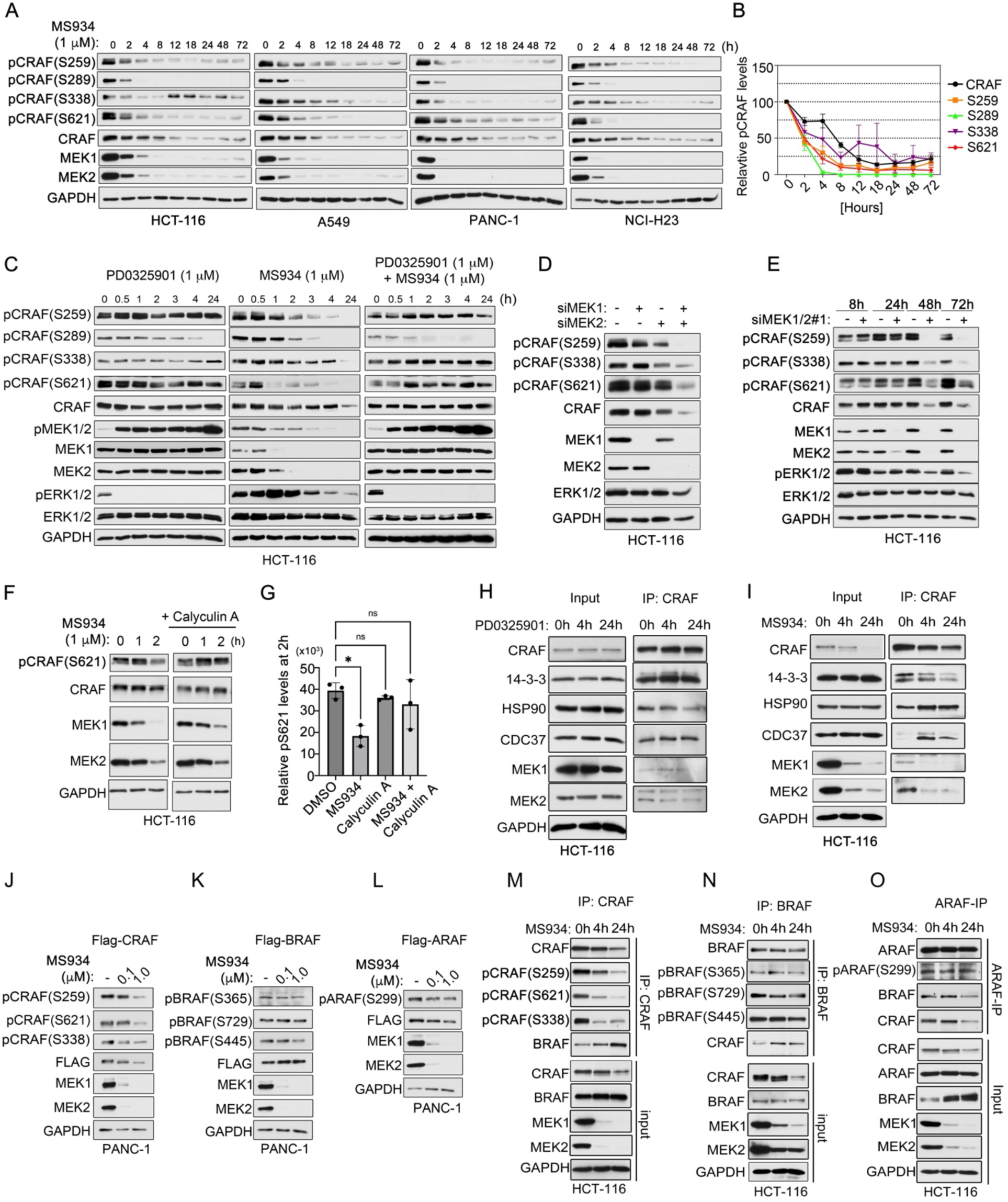
MEK1/2 degradation or knockdown caused rapid dephosphorylation of CRAF but not ARAF or BRAF, promoting protein degradation. (A-B) Immunoblot analysis of CRAF phosphorylation in HCT-116, A549, PANC-1, and NCI-H23 following 1 μM MS934 treatment across a 72-hour time course. Fresh MS934 was administered every 24 hours. Blots are representative, and densitometric analysis of CRAF total, p-S259, p- S289/296/301, p-S338, and p-S621 (B) are means ± SEM across cell line blots, each normalized to the loading control, GAPDH. CRAF, MEK1, and MEK2 blots are also shown in Fig. 2A. (C) Immunoblotting for CRAF phosphorylation in HCT-116 cells treated with 1 μM PD0325901, MS934, or the combination for 0, 0.5, 1, 2, 3, 4, or 24 hours. (D) Immunoblot analysis of CRAF phosphorylation levels in HCT-116 cells treated with non- targeting siRNAs or MEK1, MEK2, or MEK1 and MEK2 siRNAs and cultured for 48 hours. (E) Immunoblot analysis of CRAF phosphorylation levels in HCT-116 cells treated with siRNAs targeting both MEK1 and MEK2 for 8, 24, 48, and 72 hours. (F-G). Immunoblotting of CRAF phosphorylation at S621 levels in HCT-116 cells treated with 1 μM MS934 alone or in combination with 2.5 nM Calyculin A for 0, 1, or 2 hours. Cells were pre- treated with Calyculin A for 30 minutes followed by MS934 treatment for indicated times. Blots are representative, and densitometric analysis of CRAF p-S621 (G) are means ± SD of 3 independent blots, each normalized to total CRAF protein and the loading control, GAPDH. (H-I) Immunoprecipitation (IP) of endogenous CRAF from HCT-116 cells treated with 1 μM PD0325901 (H) or 1μM MS934 for 0, 4, or 24 hours. The CRAF IPs were blotted with the indicated antibodies. Input samples were used as controls. (J-L) Immunoblot analysis of RAF phosphorylation in PANC-1 cells expressing Flag-CRAF(J), Flag-BRAF (K), or Flag-ARAF (L) treated with 0, 0.1, and 1 μM MS934 for 24 hours. (M-O) Immunoprecipitation (IP) of endogenous CRAF (M), BRAF (N), or ARAF (O) from HCT-116 cells treated with 1 μM MS934 for 0, 4, or 24 hours. The RAF IPs were blotted with the indicated antibodies. Input samples were used as controls.

To determine if blocking MEK1/2 degradation using excess PD0325901 could rescue the dephosphorylation of CRAF post-MS934 treatment, we treated HCT-116 cells with 1 μM PD0325901, MS934 or the combination and monitored CRAF phosphorylation across time (**Fig. 3C**). Reductions in CRAF phosphorylation at S259, S338, and S621 were observed within 2 hours of MS934 treatment but not in response to PD0325901-treatment, whereas decreased phosphorylation of ERK1/2 negative feedback sites S289/296/301 occurred 2 hours post- PD0235901 or MS934 treatment. Notably, combined treatment of MS934 and excess PD0325901 rescued the MS934-mediated dephosphorylation of CRAF at S259, S338, and S621, but not at the ERK1/2-regulated sites S289/296/301.

Next, we examined the effect of knockdown of MEK1, MEK2, or both on CRAF phosphorylation. Genetic depletion of MEK1 or MEK2 alone did not reduce CRAF phosphorylation or total protein levels in HCT-116 cells, whereas the simultaneous knockdown of both MEK1/2 with 3 distinct siRNAs (MEK1/2-siRNA #1, MEK1/2-siRNA #2 or MEK1/2-siRNA #3) that targeted shared regions within MEK1 and MEK2 reduced CRAF phosphorylation at S259, S338 and S621 (**Fig. 3D**) as did combining MEK1 and MEK2 siRNAs (**Fig. S3B**). Additionally, immunoprecipitation of endogenous CRAF from HCT-116 cells treated with MEK1/2 siRNAs for 24 or 48 hours showed that MEK1/2 knockdown reduced CRAF phosphorylation at S259, S338, and S621 by 48 hours (**Fig. S3C**). Decreased phosphorylation of CRAF at S621, as well as total CRAF protein levels, were also observed in A549, PANC-1, and NCI-H23 cell lines following MEK1/2 knockdown (**Fig. S3D**). MEK1/2 knockdown over a 72-hour period in HCT-116 cells revealed that MEK1/2 proteins were reduced at 24 hours, whereas CRAF phosphorylation at S259, S338, and S621 was not reduced until 48 hours (**Fig. 3E**).

Phosphatases, including PP1/PP2A, dephosphorylate CRAF and other components of the MAPK signaling cascade (Boned Del Rio et al., 2019, Ory et al., 2003). To investigate whether PP1/PP2A phosphatases contributed to the dephosphorylation of CRAF by MS934, we co-treated HCT-116 or A549 cells with MS934 and the PP1/PP2A inhibitor Calyculin A for 1 or 2 hours and measured CRAF phosphorylation at S621 by immunoblot. Notably, co-treatment of MS934 and Calyculin A rescued the MS934-mediated dephosphorylation of S621 at the 2-hour time point in HCT-116 (**Fig. 3F-G)** and A549 cells (**Fig. S3E-F**), signifying that PP1/PP2A were involved in the rapid dephosphorylation of CRAF at S621 that occurred following MEK1/2 degradation.

Proper protein folding, stability, and activation of CRAF are regulated by interacting proteins 14-3-3, HSP90, and CDC37 (Thorson et al., 1998, Schulte et al., 1995, Mitra et al., 2016, Garcia-Alonso et al., 2022). Binding of 14-3-3 to CRAF phosphorylated at S259 and S621 is essential for the regulation of CRAF kinase activity (Thorson et al., 1998). To explore the effect of MEK1/2 degradation or inhibition on the interaction of CRAF with 14-3-3, CDC37, or HSP90, we performed immunoprecipitations of endogenous CRAF from HCT-116 cells treated with 1 μM PD0325901 or MS934 for 4 or 24 hours. In vehicle-treated HCT-116 cells, a protein complex of CRAF, HSP90, CDC37, 14-3-3, and MEK1/2 was observed (**Fig. 3H-I**). PD0325901 did not alter the binding of MEK1/2 to CRAF, whereas immunoprecipitation of CRAF from MS934-treated cells showed reduced MEK1/2 bound to CRAF (**Fig. 3H-I**). A reduction in 14-3-3 binding to CRAF in response to MS934 but not PD0325901-treatment at 4 and 24 hours was observed, consistent with rapid dephosphorylation of CRAF at S259 and S621, both docking sites for 14-3-3 (**Fig. 3H- I**). Conversely, an increase in CDC37 and HSP90 interaction was observed following 4 and 24 hours of treatment with MS934 but not PD0325901 (**Fig. 3H-I**), suggesting dephosphorylation of S621 caused CRAF misfolding, promoting increased chaperone activity by HSP90 and CDC37, as previously reported (Noble et al., 2008, Silverstein et al., 1998, Grammatikakis et al., 1999). Thus, although we cannot exclude the possible importance of CRAF dephosphorylation at other sites (i.e. p-S259 and p-S338), taken together these data suggest that dephosphorylation of CRAF at S621 contributed to the observed decreased stability and increased degradation of CRAF that occurred in response to MEK1/2 protein loss.

MEK1/2 degradation caused loss of CRAF but not ARAF or BRAF proteins. Next, we expressed Flag-tagged ARAF, BRAF, or CRAF in PANC-1 cells co-expressing MEK1-GFP, treated with MS934 for 24 hours and monitored phosphorylation. As expected, MS934 treatment reduced Flag-CRAF phosphorylation and total protein levels (**Fig. 3J**), but no change in phosphorylation of Flag-BRAF at S362, S446, or S729 (equivalent sites to S259, S338 and S621 in CRAF) was observed (**Fig. 3K**). MS934 treatment marginally increased Flag-BRAF total protein levels. Minimal reduction in phosphorylation of Flag-ARAF at S299 (equivalent site to S338 in CRAF) and Flag-ARAF protein levels was observed following MS934 treatment (**Fig. 3L**). Similarly, as expected immunoprecipitation of endogenous CRAF from MS934-treated HCT-116 cells showed reduced phosphorylation and total protein levels (**Fig. 3M**), whereas immunoprecipitation of BRAF (**Fig. 3N**) or ARAF (**Fig. 3O**) showed no reductions in phosphorylation or total protein levels at 4 or 24 hours post-MS934 treatment. MEK inhibition promotes CRAF-BRAF dimerization and activation (Lamba et al., 2014). Despite reduced CRAF levels, a robust increase in BRAF interaction with CRAF was observed in response to MS934 treatment like PD0325901 (**Fig. 3M-N, S3G**). Combining RAF709 with MS934 inhibited p-MEK1/2 levels more than single agents, suggesting BRAF activation occurred following MEK1/2 degradation (**Fig. S3H**). A minor reduction in ARAF binding to CRAF or BRAF was observed 24 hours post-MS934 treatment (**Fig. 3O**). Thus, MEK1/2 degradation differentially affects RAFs, causing CRAF degradation and BRAF activation.

### MEK1/2 Binding to CRAF but not Kinase Activity Required for CRAF Phosphorylation at S621 and Protein Stability

Our knockdown and degradation studies showed that MEK1/2 was required for CRAF stability and activation, therefore, we hypothesized that increasing MEK1/2 protein levels would also promote CRAF stabilization and activation. Moreover, prior studies have shown that co-expression of MEK1 and CRAF promoted increased phosphorylation of MEK1/2 and ERK1/2 (Leicht et al., 2013). Here, we showed that increasing concentrations of Myc-MEK1 and/or Myc- MEK2 caused stabilization of Flag-CRAF total protein levels, as well as elevated levels of p-S621, and p-MEK1/2 (**Fig. 4A**). In contrast, Flag-BRAF protein levels were equivalent in the presence or absence of MEK1-GFP expression, as was phosphorylation of BRAF at S729 (**Fig. S4A**). Flag- ARAF showed some increase in phosphorylation at S299 and total protein expression when co-expressed with MEK1-GFP (**Fig. S4A**), demonstrating that overexpression of MEK1/2 can also alter ARAF protein levels.

**Figure 4.**
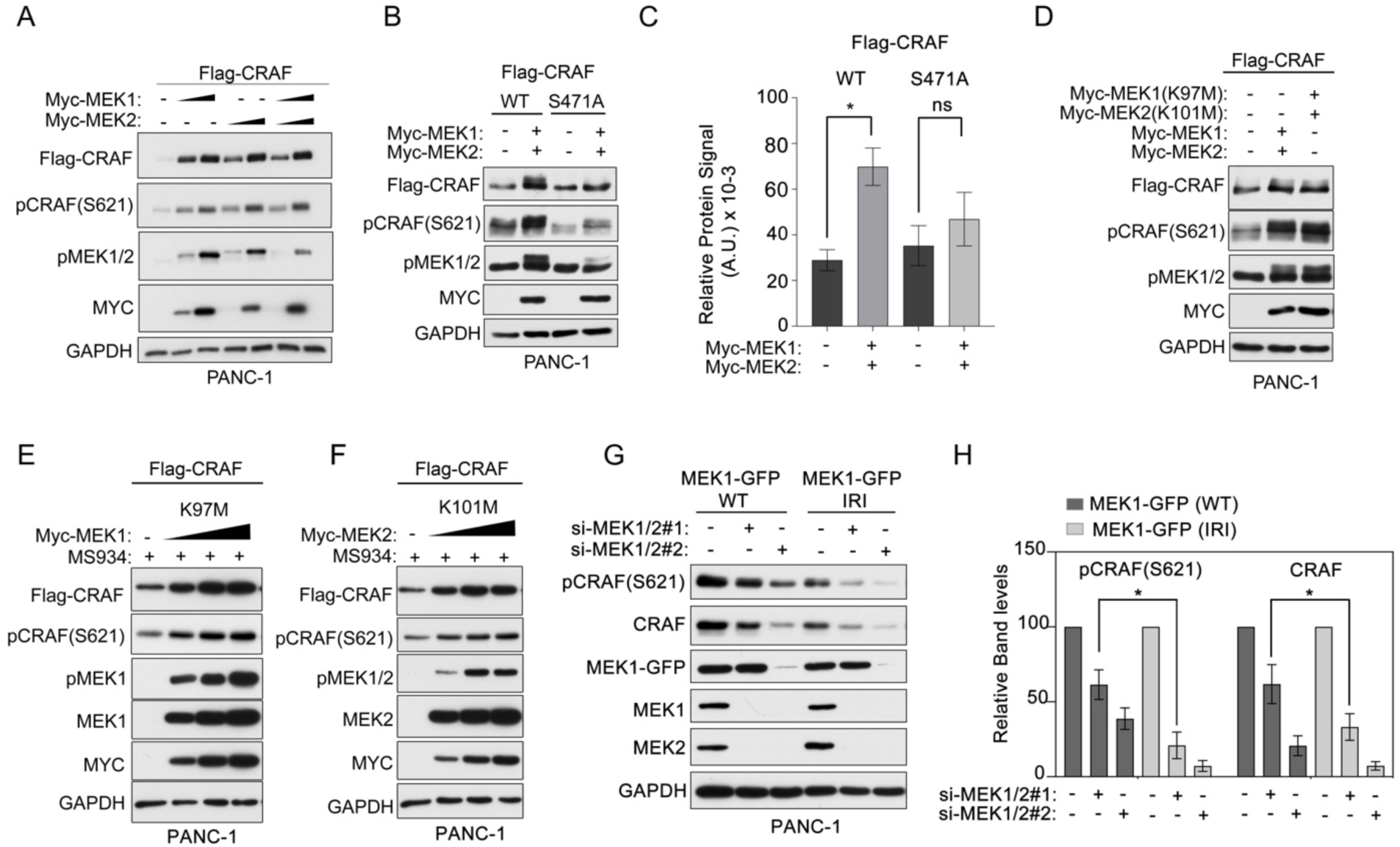
MEK1/2-binding to CRAF and not catalytic activity promotes CRAF stabilization and activation. (A) Immunoblot analysis of total and phosphorylated Flag-CRAF levels in PANC-1 cells transfected with Flag-CRAF (2.5 μg) and escalating concentrations of Myc-MEK1 (37.5, 75 ng), Myc-MEK2 (50, 100ng) or the combination of Myc-MEK1 and Myc-MEK2 (halving amounts) for 48 hrs. (B-C) Immunoblot analysis of total and phosphorylated Flag-CRAF levels in PANC-1 cells transfected with Flag-CRAF or Flag-CRAF(S471A) (2.5 μg) and Myc-MEK1 (37.5 ng)/ Myc-MEK2 (25 ng). Blots are representative, and densitometric analysis of p-S621 and CRAF (C) are means ± SD from three blots, each normalized to the loading control, GAPDH. (D) Immunoblot analysis of total and phosphorylated Flag-CRAF levels in PANC-1 cells transfected with Flag-CRAF and Myc-MEK1/2 wild-type of Myc-MEK1(K97M)/MycMEK2(K101M) (25 ng per). (E-F). Immunoblotting of total and phosphorylated Flag-CRAF levels in MS934-treated (1 μM) PANC-1 cells transfected with Flag-CRAF (2.5 μg) and escalating concentrations of Myc- MEK1(K97M) (75, 150, 300 ng) (E) or Myc-MEK2(K101M) (75, 150, 300 ng) (F) for 48 hrs. Cells were pre-treated with MS934 for 4 hours to deplete endogenous MEK1/2. (G-H) Immunoblot analysis of endogenous CRAF total and p-S621 levels in PANC-1 cells expressing si-MEK1/2#1 resistant MEK1-GFP(WT) o MEK1-GFP(IRI) treated with control, si- MEK1/2#1 or MEK1/2#2 siRNAs for 72 hours. Blots are representative, and densitometric analysis of p-S621 and CRAF (G) are means ± SEM from three blots, each normalized to the loading control, GAPDH.

Phosphorylation of CRAF at S471 is required for MEK1-binding, whereby S471A mutation impairs MEK1 binding (Zhu et al., 2005). Immunoprecipitation of Flag-CRAF(S471A) from PANC- 1 cells exhibited impaired MEK1 binding compared to Flag-CRAF wild-type whereas MEK2 is largely unaffected (**Fig. S4B**). Co-expression of Flag-CRAF(S471A) and Myc-MEK1/2 did not increase Flag-CRAF(S471) total protein or p-S621 levels as was observed for Flag-CRAF wild-type (**Fig. 4B-C**). Together, these data suggest that reducing MEK1-binding to CRAF impaired CRAF phosphorylation and protein stability.

To determine if MEK1/2 catalytic activity was required for CRAF stability, we mutated the conserved lysine in MEK1 (K97) or MEK2 (K101) rendering MEK1/2 kinase-dead (Pearson et al., 2000). The MEK1(K97M) mutant maintained its ability to interact with CRAF (**Fig. S4C**). Co-expression of Flag-CRAF and Myc-MEK1(K97M)/Myc-MEK2(K101M) increased p-S621, Flag-CRAF total protein and p-MEK1/2 like co-expression of wild-type Myc-MEK1/2 (**Fig. 4D**). The K97M and K101M MEK1/2 mutants are resistant to PD0325901 (Hatzivassiliou et al., 2013) and MS934 (**Fig. S4D**). Dose escalation of Myc-MEK1(K97M) or Myc-MEK2(K101M) in MS934- treated cells increased Flag-CRAF, p-S621 and p-MEK1/2 levels in the absence of endogenous MEK1/2 (**Fig. 4E-F**). Thus, catalytic activity of MEK1 or MEK2 was not required for promoting p- S621, protein stabilization, and activation of CRAF.

To further explore the role of MEK1 binding in regulating CRAF stability, we made alanine substitutions for Met 308 and Ile 310 in MEK1 which impairs binding to CRAF (IRI) (McKay et al., 2009). Immunoprecipitation of MEK1-GFP(IRI) from cells showed reduced binding to CRAF (**Fig. S4E**). Next, we generated MEK1/2 (#1) siRNA-resistant MEK1-GFP(IRI) mutants and evaluated the ability of the RAF binding mutant to restore endogenous CRAF p-S621 or total protein levels in HCT-116 and PANC-1 cells. Expression of siRNA-resistant MEK1-GFP(WT) partially rescued p-S621 and CRAF levels that occurred post MEK1/2 knockdown (**Fig. 4G-H, S4F-G**). Notably, expression of MEK1/2 siRNA (#1)-resistant MEK1-GFP(IRI) did not restore CRAF p-S621 or total protein levels to the extent of MEK1-GFP(WT) expression in PANC-1 or HCT-116 cells, further establishing that binding of MEK1 to CRAF was essential for promoting CRAF p-S621 and protein stability.

### MEK1/2 Binding to CRAF Essential for Maturation into Functional Kinase

Distinct cellular pools of CRAF exist in cells that include 1) newly synthesized immature CRAF and 2) mature active form of CRAF phosphorylated at S621 (Mitra et al., 2016). HSP90 inhibitors selectively degrade immature newly synthesized CRAF, preventing the formation of the active mature CRAF containing p-S621 (Mitra et al., 2016). In our characterization of MEK1/2 degradation or knockdown, we repeatedly observed a minor pool (∼25%) of CRAF remaining post-MS934 treatment (**Figs. 1C, 5A)** or MEK1/2 knockdown (**Fig. 5A**). To characterize the observed residual CRAF, we treated HCT-116 cells with MS934, the protein synthesis inhibitor, cycloheximide, the HSP90 inhibitor STA9090, or the combinations of MS934 and cycloheximide or STA9090. Notably, combining MS934 with cycloheximide enhanced the degradation of CRAF compared to single agents (**Fig. 5B**). HSP90 inhibition reduced CRAF levels like combining MS934 and cycloheximide, while combining MS934 and STA9090 did not reduce CRAF greater than HSP90 inhibition alone (**Fig. 5C**). Cycloheximide treatment also enhanced degradation of CRAF in response to MS934 treatment in A549 and HEK293T (**Fig. S5A-B**). Additionally, combined blockade of translation using MTOR inhibitor RAD001 in MS934-treated HCT-116 cells (**Fig. 5D**) or Torin-1 in MS934-treated A549 cells (**Fig S5C**) enhanced the depletion of p-S621 and total CRAF levels. Together, these findings suggested that the minor pool of CRAF remaining post-MEK1/2 degradation was newly synthesized immature CRAF that was sensitive to protein synthesis or MTOR inhibition.

**Figure 5.**
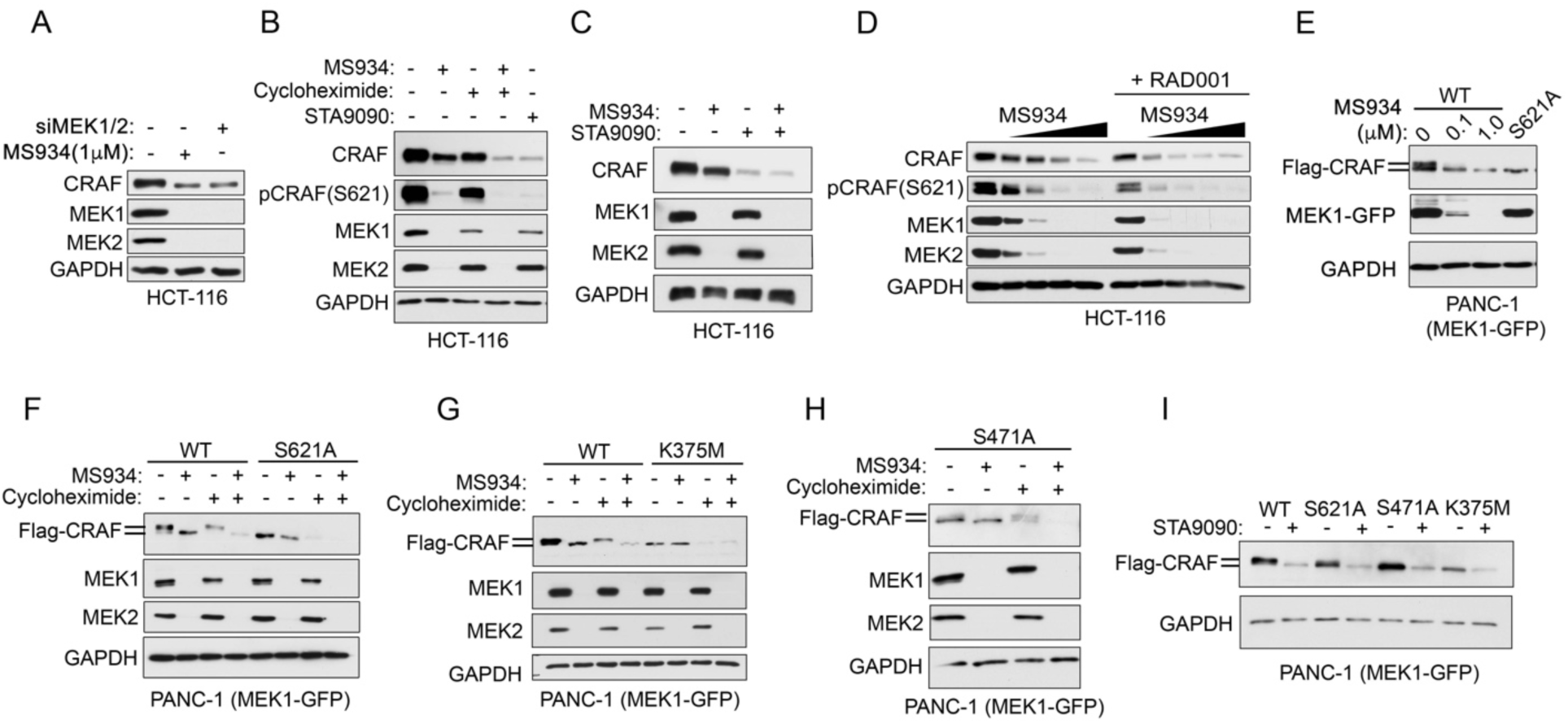
MEK1/2 is required for stabilizing mature CRAF but not the immature form. (A) Immunoblot analysis of CRAF protein levels in HCT-116 cells treated with MEK1/2 siRNAs for 72 hours or 1 μM MS934 for 24 hours. (B) Immunoblotting for CRAF total and p-S621 levels in HCT-116 cells treated with DMSO, 1 μM MS934, 100 μg/ml cycloheximide, 0.05 μM STA9090, or the combination of MS934 and cycloheximide. (C) Immunoblot analysis of RAF levels in HCT-116 cells treated with DMSO, 1 μM MS934, 0.05 μM STA9090, or the combination for 24 h. (D) Immunoblot analysis for p-S621 and CRAF levels in HCT-116 cells treated with DMSO or escalating doses of MS934 in the presence or absence of 2 μM RAD001 for 24 hours. (E) Immunoblotting for Flag-CRAF levels in PANC-1 cells transfected with Flag-CRAF(WT) or Flag-CRAF(S621A) and cells expressing Flag-CRAF(WT) were treated with DMSO, 0.1 or 1 μM MS934. (F-H) Immunoblot analysis of Flag-CRAF levels in PANC-1 cells stably expressing MEK1-GFP transfected with Flag-CRAF(WT) or Flag-CRAF(S621A) (F) or Flag-CRAF(WT) or Flag- CRAF(K375M) (G) or Flag-CRAF(S471A) treated with DMSO, 1 μM MS934, 100 μg/ml cycloheximide, or the combination of MS934 and cycloheximide for 24 hours. (I) Immunoblotting for Flag-CRAF levels in PANC-1 cells stably expressing MEK1-GFP transfected with Flag-CRAF(WT), Flag-CRAF(S621A), Flag-CRAF(S471A) or Flag- CRAF(K375M) treated with DMSO, or 0.05 μM STA9090 for 24 hours.

Exogenous expression of CRAF presents as two bands in immunoblots, the upper band being the phosphorylated form of CRAF that contained p-S621 amongst other phosphosites (Ferrier et al., 1997). Notably, MS934 treatment reduced the upper Flag-CRAF band, while some residual lower Flag-CRAF band remained that migrated like the immature Flag-CRAF(S621A) mutant (**Fig. 5E**). To explore the stability of mature and immature pools of CRAF, we treated PANC-1(MEK1-GFP) cells co-expressing maturation-competent Flag-CRAF(WT), or maturation- noncompetent Flag-CRAF(S621A), or kinase-dead Flag-CRAF(K375M) mutant with 1 μM MS934, cycloheximide, or the combination. MS934 treatment reduced the upper and partially the lower Flag-CRAF(WT) band, while cycloheximide reduced the lower Flag-CRAF(WT) band, sparing the upper phosphorylated band (**Fig. 5F-G**). Notably, the combination of MS934 and cycloheximide reduced both Flag-CRAF(WT) upper and lower bands to a greater extent than single agents, confirming that the lower band observed post-MS934 was newly synthesized Flag-CRAF (**Fig. 5F-G**). Additionally, whereas MS934 treatment reduced the upper mature FLAG-CRAF(WT) band, it did not reduce immature Flag-CRAF(S621A) or Flag-CRAF(K375M) levels (**Fig. S5D-E**). In contrast, cycloheximide treatment depleted the majority of Flag-CRAF(S621A) or Flag-CRAF(K375M) levels, and the combination of MS934 and cycloheximide was minimally better than cycloheximide treatment alone (**Fig. 5F-G**). Expression of the Flag-CRAF(S471A) MEK1-binding mutant also presented as predominantly a single lower band that was insensitive to MS934 like the CRAF(K375M) or CRAF(S621A) mutants (**Fig. 5H, S5F-G**). Depletion of Flag-CRAF(S471A) levels was detected with cycloheximide alone, demonstrating that in the absence of MEK1-binding, CRAF(S471A) predominantly existed as the immature form of CRAF. As expected, HSP90 inhibition reduced both forms of Flag-CRAF(WT) and immature Flag-CRAF(S621A), Flag-CRAF(K375M) or FLAG-CRAF(S471A) mutant protein levels (**Fig. 5I**).

Together, these findings demonstrated that CRAF mutants unable to autophosphorylate at S621 or that exhibit impaired MEK1-binding were insensitive to MS934, predominately existed as the immature form of CRAF, and were more sensitive to protein synthesis inhibition than mature CRAF.

### Immature CRAF Has Anti-Apoptotic Functions Independent of MEK1/2

CRAF has been shown to have kinase-independent functions outside the MEK1/2 signaling axis (McCormick, 2018). CRAF promotes cell survival by inhibiting apoptosis through a kinase-independent mechanism involving binding to apoptosis signal-regulating kinase 1 (ASK1) (Chen et al., 2001). Notably, CRAF knockdown by shRNA in HCT-116 and A549 cells caused an increase in cleaved PARP, indicating activation of caspase activity and apoptosis **(Fig. 6A-B**). A time course analysis of CRAF knockdown in A549 cells showed that CRAF protein levels were sufficiently reduced to promote cleaved PARP by 72 hours post-doxycycline induction (**Fig. S6A**). In contrast, depletion of the mature form of CRAF and MEK1/2 by 1 μM MS934 treatment only minimally increased cleaved PARP levels compared to CRAF knockdown **(Fig. 6A-B**). Strikingly, knockdown of the residual immature CRAF post-MS934 treatment in HCT116 or A549 cells by shRNA induced apoptosis by 48 hours (**Fig. 6C-D**), suggesting that the inability of MS934 to induce apoptosis could be due to insufficient blockade of CRAF anti-apoptotic functions outside the MEK-ERK pathway. Consistent with this, immunoprecipitation of Flag-CRAF in MS934- treated PANC-1(MEK1-GFP) cells revealed an increased interaction between ASK1 and residual CRAF compared to vehicle treatment (**Fig. 6E**). A similar phenomenon was observed with immature kinase dead Flag-CRAF(K375M) (**Fig. 6E**). Together, these findings demonstrated that post-MEK1/2 degradation the residual immature CRAF promoted survival outside the MEK1/2 pathway via binding and antagonizing ASK1.

**Figure 6.**
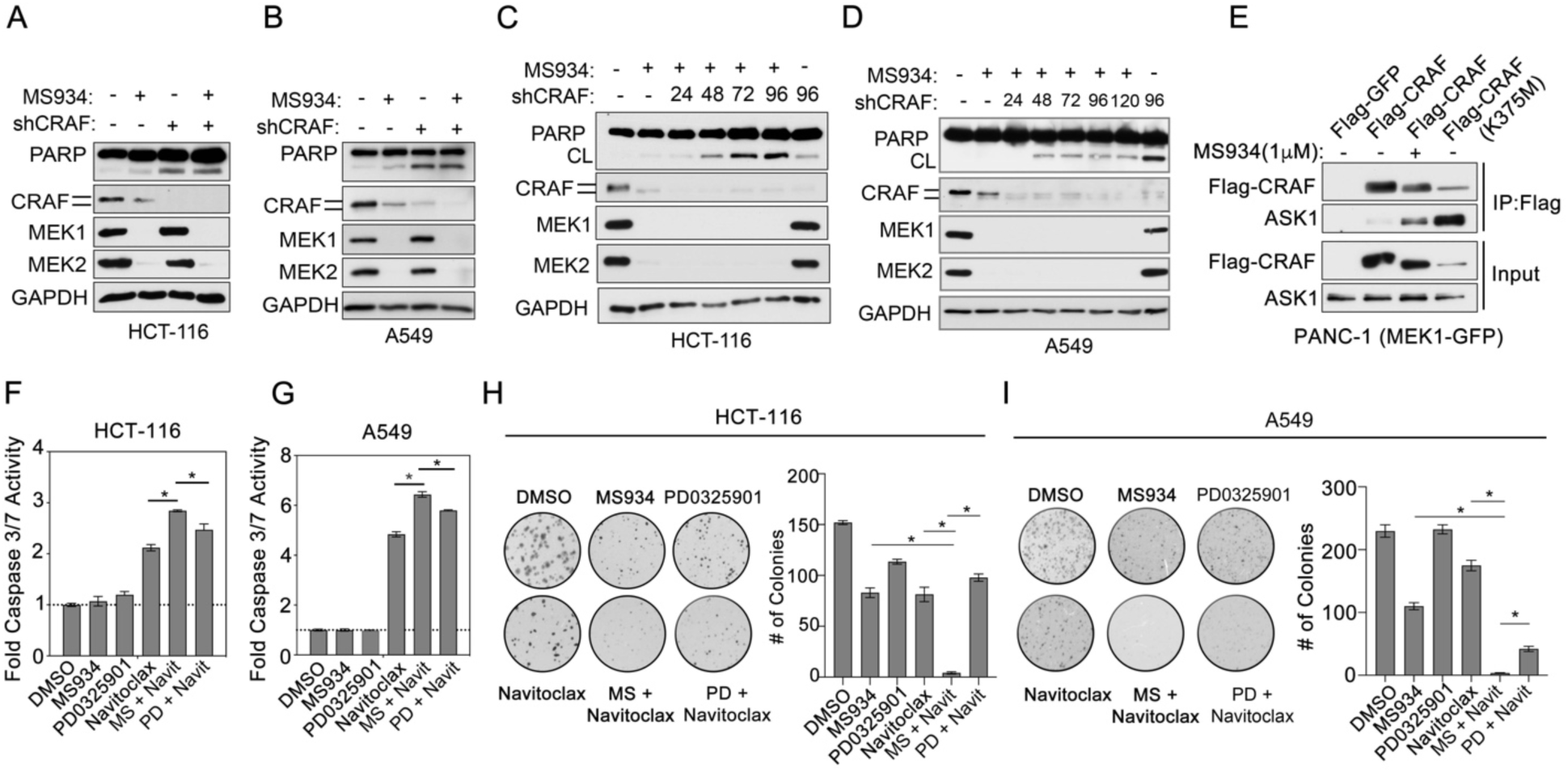
Immature CRAF has anti-apoptotic functions outside the MAPK signaling pathway. (A-B) Immunoblot analysis of cleaved PARP levels in HCT-116 (A) or A549 (B) cells treated with DMSO or 1 μM MS934 or upon short hairpin RNA (shRNA)-mediated depletion of CRAF (shCRAF) or uninduced (1 μg/ml doxycycline for 120 h (A), 72 h (B). (C-D) Immunoblot analysis of cleaved PARP levels in HCT-116 (C) or A549 (D) cells treated with DMSO or 1 μM MS934 for 24 h followed by short hairpin RNA (shRNA)-mediated depletion of CRAF (shCRAF) or uninduced (1 μg/ml doxycycline for 96 h (C) or 120 h (D)). (E) Immunoprecipitation (IP) of Flag-CRAF from PANC-1 cells stably expressing MEK1-GFP treated with DMSO or 1 μM MS934 for 24 h or IP of Flag-CRAF(K375M). The Flag-CRAF IPs were blotted with the indicated antibodies. IP of Flag-GFP and input samples were used as controls. (F-G) Caspase 3/7 activation analysis in HCT-116 (F) or A549 (G) cells treated with DMSO, 0.005 μM PD0325901, 0.5 μM MS934, 1 μM Navitoclax, or the combination of PD0325901 or MS934 and Navitoclax for 24 h. Data are means ± SEM of three independent experiments. (H-I) Soft-agar colony formation assay in HCT-116 (H) or A549 (I) cells treated with DMSO, 0.005 μM PD0325901, 0.5 μM MS934, 1 μM Navitoclax, or the combination of PD0325901 or MS934 and Navitoclax for 2 weeks. Images are magnifications of representative fields. The corresponding colony counts are means ± SEM derived from multiple fields (N=6).

An ASK1/Jun N-terminal protein kinase pathway phosphorylates *B-cell lymphoma 2* (BCL2) inactivating its anti-apoptotic function promoting cell death (Yamamoto et al., 1999). As immature CRAF blocks apoptosis via inhibiting ASK1, we hypothesized that inhibition of BCL2 in MS934-treated cells would enhance caspase activation. Indeed, treatment of HCT-116 or A549 cells with the combination of the BCL2 inhibitor, Navitoclax, and MS934 enhanced caspase-3/7 activation compared to single agent treatments (**Fig. 6F-G**). Notably, blocking BCL2 function in MS934-treated cells was more effective at activating caspase 3/7 than combining Navitoclax and PD0325901. Moreover, the combination of Navitoclax and MS934 blocked cell viability (**Fig. S6B- C**) and anchorage-independent colony growth of HCT-116 and/or A549 cells to a greater extent than single agent treatments or the combination of Navitoclax and PD0325901 (**Fig. 6H-I**).

### A Stable Complex of CRAF-MEK1/2 Exists That is Primed for Activation by RAS Binding and RAF Dimerization

To define when MEK1/2 binding to CRAF occurs in the process of CRAF maturation, we expressed various Flag-CRAF mutants in PANC-1(MEK1-GFP) or HEK293T cells, immunoprecipitated Flag, and probed for MEK1 and MEK2 interactions. Both MEK1-GFP and endogenous MEK1 interacted with Flag-CRAF(WT), Flag-CRAF(R89L) (impaired RAS binding), Flag-CRAF(R401H) (impaired dimerization), Flag-CRAF(S621A) and Flag-CRAF(S259A/S621A), whereas the Flag-CRAF(S471A) and Flag-CRAF(K375M) mutants showed reduced binding to MEK1 (**Fig. 7A, S7A-B**). Endogenous MEK2 interacted with all Flag-CRAF mutants, suggesting there may be differences in how MEK1 and MEK2 interact with CRAF (**Fig. 7A, S7A-B**). Notably, pulldown of MEK1/2 with the Flag-CRAF(S259A/S621A) or Flag-CRAF(S621A) mutants revealed that MEK1/2 binds to CRAF independent of p-S621 status. Additionally, we observed that HSP90 and CDC37 interacted with all Flag-CRAF constructs, while 14-3-3 only interacted with Flag-CRAF(WT), Flag-CRAF(R89L), Flag-CRAF(R401H) and minimally with Flag-CRAF(S471A) (**Fig. 7A**). We detected BRAF bound to Flag-CRAF(WT) and weakly to Flag-CRAF(R89L) with minimal BRAF interaction observed with the other CRAF mutants (**Fig. 7A**), demonstrating some CRAF-BRAF dimerization occurs in the absence of CRAF-based RAS binding.

**Figure 7.**
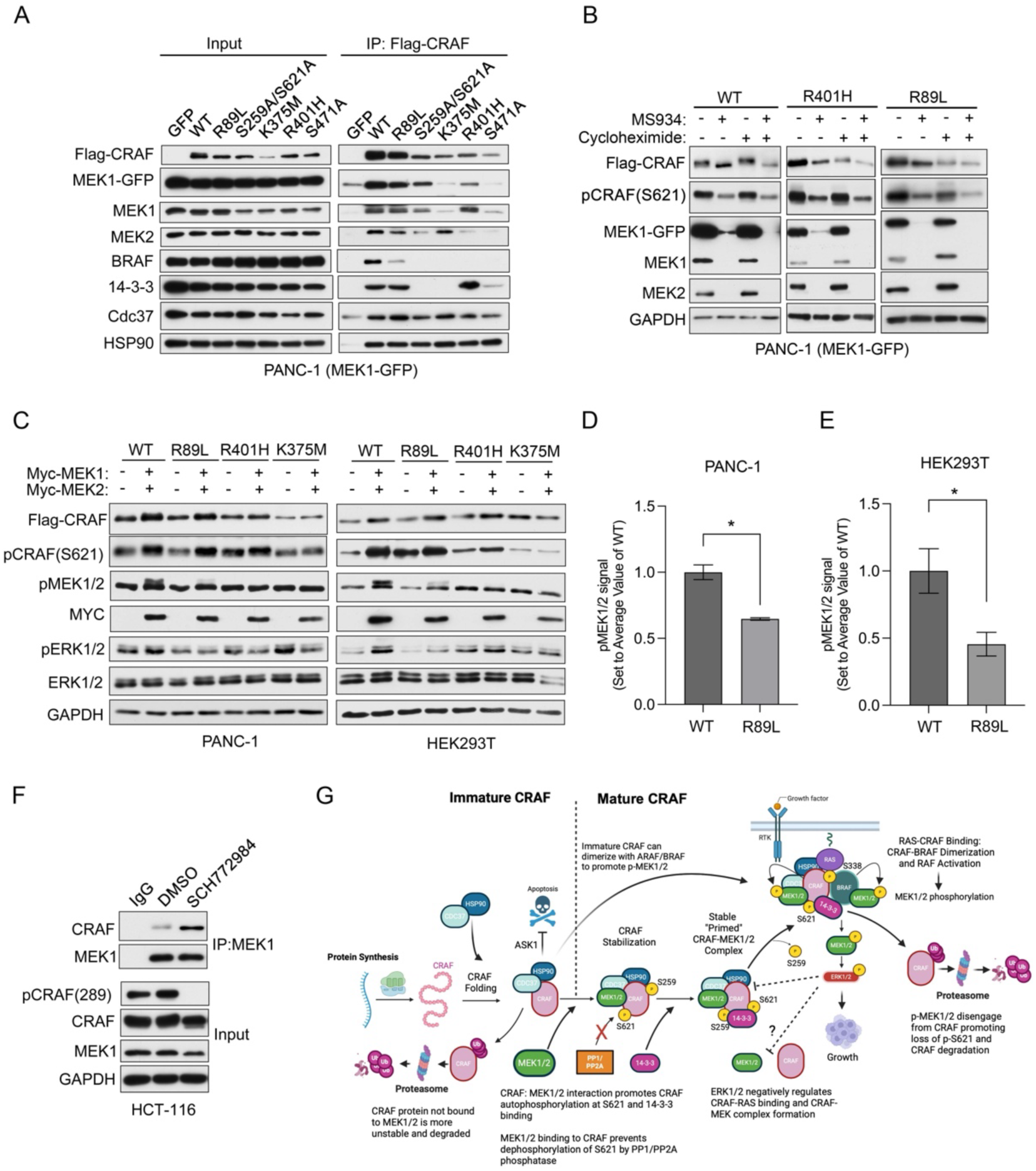
A stable complex of CRAF-MEK1/2-HSP90-14-3-3 exists that is primed for activation by RAS. (A) Immunoprecipitation (IP) of Flag-tagged CRAF constructs from PANC-1 cells stably expressing MEK1-GFP. The Flag-CRAF IPs were blotted with the indicated antibodies. IP of Flag-GFP and input samples were used as controls. (B) Immunoblot analysis of Flag-tagged CRAF constructs in PANC-1 cells stably expressing MEK1-GFP transfected with Flag-CRAF(WT), Flag-CRAF(R401H) or Flag-CRAF(R89L) treated with DMSO, 1 μM MS934, 100 μg/ml cycloheximide, or the combination of MS934 and cycloheximide for 24 hours. (C-E) Immunoblot analysis of CRAF signaling in PANC-1 cells co-transfected with Myc- MEK1/Myc-MEK2 and Flag-CRAF(WT), Flag-CRAF(R89L), Flag-CRAF(R401H) or Flag-CRAF(K375M) for 48 hrs. Blots are representative, and densitometric analysis of p-MEK1/2 and (D/E) are means ± SEM from three blots, each normalized to the loading control, GAPDH, Myc- tag and Flag levels. (F) Immunoprecipitation (IP) of endogenous MEK1 from HCT-116 cells treated with 1 μM ERK1/2 inhibitor SCH772984 for 24 hours. The MEK1 IP was blotted for CRAF and MEK1 proteins. Blots are representative. (G) Model of MEK1/2 regulation of CRAF stabilization, maturation, and activation. Figure created in BioRender. Duncan, J. (2025) https://BioRender.com/t50o519.

To determine if MEK1/2 binding to Flag-CRAF(R89L), or Flag-CRAF(R401H) preserved p-S621 and protein stability like wild-type, we treated cells with MS934, cycloheximide, or the combination and measured Flag-CRAF and p-S621 levels. MS934 treatment reduced p-S621 and total protein levels of Flag-RAF(R89L) or Flag-CRAF(R401H), and the combination of MS934 and cycloheximide enhanced degradation of the Flag-CRAF(R89L) or Flag-CRAF(R401H) mutants as observed for Flag-CRAF(WT) (**Fig. 7B**). Together, our MEK1/2 data suggest that 1) MEK1/2 interaction with CRAF occurs before RAS binding and/or RAF dimerization to promote phosphorylation of p-S621 and stabilization of total CRAF protein levels and 2) a primed complex of CRAF-MEK1/2 (including HSP90-CDC37-14-3-3) forms in cells independent of RAF dimerization and before RAS interaction.

Next, we assessed whether RAS binding or RAF dimerization was required to activate the CRAF-MEK1/2 complex. Co-expression of Myc-MEK1/2 and Flag-CRAF(R89L) or Flag-CRAF (R401H) in PANC-1 or HEK293T cells increased Flag-CRAF protein and p-S621 levels like Flag-CRAF(WT), whereas no change in Flag-CRAF protein or p-S621 was observed with the kinase-dead Flag-CRAF(K375M) mutant (**Fig. 7C**). As expected, expression of Flag-CRAF(WT) with Myc-MEK1/2 increased p-MEK1/2 in PANC-1 and HEK293T cells. Increased levels of p-MEK1/2 were also detected following expression of the Flag-CRAF(R89L) mutant with Myc-MEK1/2, consistent with prior studies demonstrating that MEK1 activates CRAF through a RAS-independent mechanism (Leicht et al., 2013). Notably, the phosphorylation of MEK1/2 by the Flag-CRAF(WT) was significantly greater than observed for the Flag-CRAF(R89L) mutant in PANC-1 and HEK293T cells (**Fig. 7D-E**). No change in p-MEK1/2 levels was observed with the Flag-CRAF(R401H) resembling the kinase-dead Flag-CRAF(K375M) mutant. Similarly, expression of Flag-CRAF(R89L) or Flag-CRAF(R401H) in PANC-1(MEK1-GFP) cells did not activate MEK or ERK like Flag-CRAF(WT) (**Fig. S7C**). Together, these findings showed that the CRAF-MEK1/2 complex required RAS interaction and RAF dimerization for optimal activation of CRAF and subsequent phosphorylation of MEK1/2.

Phosphorylation of CRAF by ERK1/2 negatively regulates the interaction between CRAF and RAS, controlling CRAF activity (Dougherty et al., 2005). MEK inhibition with inhibitors such as PD0325901 increases CRAF and MEK1 interactions due to alleviation of ERK1/2 negative feedback on CRAF (Lito et al., 2014). Here, we showed that directly inhibiting ERK1/2 activity using SCH772984 in HCT-116 cells reduced ERK-regulated sites p-S289/296/301 promoting increased interaction between MEK1 and CRAF (**Fig. 7F**). These findings suggest that ERK1/2 activity could control formation of a primed CRAF-MEK complex, competent to create a RAS-RAF-MEK signalosome.

Collectively, our findings support a revised model of CRAF activation (**Fig. 7G**) in which MEK1/2 binding to newly synthesized immature CRAF, along with HSP90-CDC37, promotes phosphorylation at S621, and protein stabilization. A CRAF-MEK-HSP90-CDC37-14-3-3 complex forms, primed for activation by RAS binding. Upon RAS engagement, CRAF dimerizes with BRAF, forming an active signalosome whereby RAFs phosphorylate MEK, leading to ERK activation and promoting cell growth. Once phosphorylated, MEK1/2 could dissociate from CRAF promoting dephosphorylation of S621 and subsequent CRAF degradation. Binding of the CRAF-MEK complexes to RAS, as well as CRAF-MEK complex formation, are negatively regulated by active ERK1/2. Finally, although the majority of CRAF in cells is the mature form, a minor pool of immature CRAF exists in cells that has anti-apoptotic functions and can also dimerize with ARAF/BRAF to promote MAPK signaling.

## DISCUSSION

Kinases are largely defined by their catalytic function despite many possessing roles independent of their kinase activity (Rauch et al., 2011). Regarding MEK1/2, the canonical function is the phosphorylation of ERK1/2 in the MAPK cascade. Such roles have been confirmed using kinase inhibitors, which only assess catalytic function. Not surprisingly, many enzymes, including kinases, are involved in structured protein complexes but their involvement in the stability of the complex remains understudied. Here, using MEK1/2 PROTACs, we identified a new kinase-independent function of MEK1/2 whereby MEK1/2 regulate the stability of the upstream kinase CRAF, but not the related ARAF or BRAF proteins. MEK1/2 facilitate the maturation of CRAF to an active kinase allowing initiation of the MAPK cascade via dimerization with ARAF/BRAF in a RAS-dependent manner. Without MEK1/2, a pool of immature CRAF maintains its anti-apoptotic functions as previously reported.

CRAF is comprised of two cellular pools, an immature form devoid of kinase activity, and the mature active form harboring p-S621 that potentiates the MAPK signaling cascade (Mitra et al., 2016). The maturation of CRAF is orchestrated by chaperones CDC37 and HSP90, which ensure proper folding of newly synthesized CRAF rendering the kinase capable to induce autophosphorylation of S621 and subsequent maturation (Silverstein et al., 1998, Grammatikakis et al., 1999, Smith et al., 2009, Mitra et al., 2016, Garcia-Alonso et al., 2022). HSP90 inhibitors selectively target immature CRAF causing its degradation, while exhibiting no effect on mature CRAF. In contrast, we discovered MEK1/2 degradation caused preferential loss of mature active CRAF. These findings illustrate the importance of both HSP90 and MEK1/2 in CRAF maturation. Interestingly, both MEK1/2 and HSP90-CDC37 bind the immature Flag-CRAF(S259A/S621A) mutants, demonstrating that MEK1/2 can associate with CRAF independent of S621 phosphorylation status. HSP90 inhibition or MEK1/2 degradation caused loss of CRAF p-S621 levels, suggesting MEK1/2 binding and HSP90 activity are both required for CRAF autophosphorylation at S621. Moreover, the binding of HSP90-CDC37 to residual CRAF devoid of S621 phosphorylation post-MS934 treatment and the impaired MEK1/2 binding CRAF(S471A) mutant indicate that HSP90-CDC37 chaperone activity is not sufficient to promote CRAF autophosphorylation at S621 in the absence of MEK1/2 binding.

Bystander proteins interacting with PROTAC targets have been reported to be collaterally degraded due to being in proximity of the recruited E3 ubiquitin ligase and proteasome machinery (Xiong et al., 2022). A detailed temporal evaluation of CRAF ubiquitination and degradation revealed that CRAF was degraded via a conserved cell-intrinsic mechanism rather than collateral degradation. The E2/E3 ligases responsible for promoting the degradation of CRAF in response to MEK1/2 degradation have not been identified, but if they were, could facilitate the development of CRAF degraders or molecular glues. It is likely that there are no published CRAF degraders because most RAF inhibitors target the active form of CRAF (Singh et al., 2023, Shao et al., 2018). Degraders targeting immature CRAF will be most advantageous, as these agents will deplete both immature and mature forms of CRAF, leading to apoptosis. HSP90 inhibitors promote degradation of the immature form of CRAF but are not selective given that HSP90 has hundreds of client proteins (Kamal et al., 2004). Alternatively, knockdown of the chaperone CDC37 promotes CRAF protein loss (Smith et al., 2009). Structural investigation of the unique CDC37-CRAF interface demonstrates that its disruption affects CRAF protein stability, highlighting the need for inhibitors or “busters” of this interaction that would selectively target immature CRAF, sparing other HSP90-CDC37 client proteins (Garcia-Alonso et al., 2022).

The current CRAF activation cycle places MEK1/2 recruitment following RAS binding and CRAF dimerization (Lavoie and Therrien, 2015). Here, we demonstrate MEK1/2 binding to CRAF occurs before RAS engagement and RAF dimerization, whereby MEK1/2 kinase-independent functions promote CRAF protein phosphorylation, stabilization, and maturation. Upon MEK1/2 depletion, CRAF is rapidly dephosphorylated at S621 by PP1/2 phosphatases, impairing 14-3-3 binding to CRAF, promoting its degradation. Prior studies showed that MEK1 activates CRAF through a RAS-independent mechanism that required MEK1-binding (Leicht et al., 2013). Here, we find the CRAF(R89L) RAS binding mutant can promote some MEK1/2 and ERK1/2 phosphorylation, though substantially less than that observed with wild-type CRAF. Notably, immunoprecipitation of the Flag-CRAF(R89L) mutant detected interaction with BRAF, suggesting that activation of the MAPK pathway by the Flag-CRAF(R89L) could involve dimerization of a preformed CRAF-MEK complex with BRAF. This is likely, as co-expressing the CRAF(R401H) dimerization mutant with MEK1/2 caused minimal phosphorylation of MEK, signifying that dimerization of CRAF was essential to achieve MEK1/2 phosphorylation. We cannot exclude the possibility that dimerization of immature CRAF with BRAF could substitute for MEK chaperone function in forming an active signalosome. Thus, our current findings suggest MEK1/2 binding to HSP90-CDC37-bound CRAF promotes CRAF protein stabilization and maturation, and that a primed complex of CRAF-MEK is formed that requires RAS binding and RAF dimerization to form a fully active signalosome to achieve optimal phosphorylation of MEK1/2.

Direct negative feedback phosphorylation of CRAF at sites within its RAS binding domain by ERK1/2 releases CRAF from RAS, dissociating RAF dimers and inhibiting the MAPK cascade, with dephosphorylation of CRAF by PP2A and PIN1 promoting MAPK reactivation (Dougherty et al., 2005). Here, we discovered binding of MEK1/2 to CRAF promotes stabilization and maturation, but how the recruitment of MEK to CRAF is regulated to promote CRAF activation remains undetermined. MEK1/2 blockade by inhibitors (i.e., PD0325901) relieves ERK-mediated feedback inhibition of CRAF promoting CRAF-MEK protein interaction (Lito et al., 2014), suggesting ERK1/2 activity may directly regulate MEK1/2-CRAF complex formation. Consistent with this, we showed the ERK1/2 inhibitor SCH772984 promoted an association of CRAF with MEK1. These findings suggest that activated ERK1/2 may negatively regulate the interaction of CRAF and MEK1, and that once ERK1/2 is inactivated by dephosphorylation, MEK1/2 binds to CRAF promoting maturation and activation of CRAF in a new primed complex. Interestingly, stoichiometric analysis of MEK1 and CRAF levels in cells showed that there is 100-fold more MEK than CRAF in cells (Fujioka et al., 2006); yet CRAF activity is tightly controlled. Notably, proteins such as BRAF or KSR can sequester MEK from CRAF by dimerizing kinase domain to kinase domain; similarly MEK-MEK dimerization (Catalanotti et al., 2009) could limit CRAF-MEK complex formation in quiescent cells (Haling et al., 2014, Catalanotti et al., 2009).

Despite similar structures and functions, CRAF and BRAF exhibit opposing behaviors in response to MEK1/2 degradation, (i.e. dephosphorylation and degradation of CRAF and activation of BRAF). Mutation of the S621 autophosphorylation site to an alanine or kinase inactivating mutations renders CRAF proteolytically unstable resulting in increased degradation (Noble et al., 2008). In contrast, kinase-dead BRAF did not impact BRAF protein stability, signifying BRAF and CRAF stability and activation are differentially regulated in cells (Noble et al., 2008). Notably, BRAF harbors evolutionarily conserved aspartic residues in its N-terminal region that mimic phosphorylation producing a constitutively active kinase, whereas CRAF requires phosphorylation at several sites to become an active kinase (Mason et al., 1999). BRAF also does not require the complex orchestration of chaperone activity associated with CRAF as demonstrated by HSP90 inhibition (Grbovic et al., 2006). Like MEK1/2 inhibition (Lito et al., 2014), we found degradation of MEK1/2 promoted BRAF dimerization with residual immature CRAF activating BRAF, signifying that immature CRAF, devoid of kinase activity, can promote MAPK signaling. Our findings are consistent with studies that showed CRAF dimerization rather than activity promotes KRAS tumorigenesis (Venkatanarayan et al., 2022). Activation of BRAF by MS934 will likely be a resistance mechanism, requiring co-inhibition of BRAF, with agents like RAF709, with MEK1/2 degraders to sustain suppression of MEK-ERK activation. BRAF and MEK1 form an inactive complex that is recruited to the membrane upon RAS activation, promoting BRAF dimerization and activation (Haling et al., 2014). Without MEK1/2 proteins, BRAF is free from MEK1 regulation and able to dimerize with immature CRAF promoting BRAF activation. Similar findings regarding BRAF activation showed that MEK1 binding to KSR1 promotes BRAF- KSR1 dimerization independent of MEK1 kinase activity (Lavoie et al., 2018). Thus, MEK1/2 binding to BRAF may negatively regulate BRAF dimerization with CRAF, controlling BRAF activation. In contrast, MEK1/2 binding dictates CRAF stability and activation.

MS934 promotes the degradation of both MEK1 and MEK2, causing subsequent loss of CRAF in cells. Genetic knockdown of endogenous MEK1 or MEK2 alone did not cause CRAF degradation, demonstrating that both MEK1 and MEK2 binding to CRAF are required for CRAF stabilization, phosphorylation, and maturation. Interestingly, we demonstrate that MEK1 and MEK2 have differential binding preferences for CRAF mutants, finding MEK2, but not MEK1, bound to CRAF(S471A) and CRAF(K375M). Mutation of lysine 375 blocks interaction with MEK1 in assays with purified proteins, demonstrating CRAF ATP-binding was necessary for MEK1/2 binding (Xiang et al., 2002). MEK1 binding to CRAF is likely kinase domain to kinase domain, whereby disruption of the catalytic lysine in CRAF could disrupt kinase domain interactions. How the S471A mutation impairs MEK1 binding to CRAF has not been determined. The fact that MEK2 can bind the CRAF(K375M) or CRAF(S471A) mutants suggests MEK2 may interact with a region of CRAF distinct from MEK1, allowing the possibility for simultaneous binding of MEK1 and MEK2 to wild-type CRAF. Further molecular and structural studies defining the MEK1 and MEK2 binding sites within CRAF, and whether MEK2 functions differently than MEK1 in the stabilization and maturation of CRAF will be of particular interest. MEK2 has a unique role in apoptosis (Huth et al., 2016), while MEK1 serves as the predominant MEK in RAS-driven MAPK signaling (Chen et al., 2001), suggesting MEK2 may regulate the anti-apoptotic function of immature CRAF.

CRAF has both kinase-dependent and kinase-independent functions through activation of MEK-ERK signaling and promoting growth and survival of KRAS mutant cancers, respectively (McCormick, 2018). Genetic depletion of CRAF but not its inhibition leads to growth repression and apoptosis in KRAS mutant cancers (Sanclemente et al., 2021, Venkatanarayan et al., 2022). Kinase-independent functions of CRAF suppress apoptosis in KRAS mutant cells via interaction with pro-apoptotic kinases ASK1 and MST2, inhibiting their kinase activities (O’Neill et al., 2004, Chen et al., 2001) Consistent with these findings, we showed that without MEK1/2 present the immature form of CRAF exhibits anti-apoptotic functions via ASK1 and BCL2 (Yamamoto et al., 1999). Moreover, we demonstrate that MEK1/2 degradation promotes dependency on immature CRAF’s anti-apoptotic function, which sensitizes MS934-treated cells to BCL2 inhibition. It will be of particular interest to determine whether the anti-apoptotic functions of residual immature CRAF will promote resistance and disease persistence following chronic MS934 exposure. Additionally, deconvoluting interacting partners of immature CRAF and mature CRAF could reveal novel functions of immature CRAF outside of MEK-ERK signaling. Kinase-independent functions of CRAF have been implicated in DNA damage (Advani et al., 2015) and cell cycle progression (Mielgo et al., 2011). Whether these roles require MEK1/2 binding or are orchestrated by immature CRAF remains to be determined.

CRAF degradation represents a highly advantageous therapeutic target for the treatment of MAPK-altered cancers; however, there are no published CRAF degraders. Our findings suggest that developing CRAF degraders may prove difficult using existing RAF inhibitors due to the complex maturation process of CRAF. Here, we identified an alternative strategy to deplete the majority of CRAF protein in cells by exploiting the kinase-independent function of MEK1/2 in preserving CRAF protein stability. Our findings support the use of MEK1/2 degraders combined with translation inhibitors, such as MTOR inhibitors, to degrade both immature and mature CRAF or targeting the anti-apoptotic functions of immature CRAF using BCL2 inhibitors. Further studies are needed to determine the effectiveness of such therapies in animal models, as well as the potential safety issues related to systemic depletion of CRAF and MEK1/2 proteins. Notably, in prior work, we showed that MS934 treatment was safely administered *in vivo* and blocked tumor growth when combined with EGFR inhibitors in KRAS mutant colorectal cancer xenografts (Kurimchak et al., 2022).

## EXPERIMENTAL PROCEDURES

### Cell Lines

Cell lines were verified by IDEXX laboratories and free of mycoplasma. All media with the exception of the Mammary Epithelial Cell Basal Medium was supplemented with 10% FBS, 100 U/ml Penicillin-Streptomycin and 2mM GlutaMAX. HCT-116, NCI-H23, A549, HT-29, and OVCAR8 cell lines were maintained in RPMI-1640. HEK293T cells were maintained in DMEM. PANC-1 and HPAF-II cells were maintained in DMEM supplemented with 1mM Sodium Pyruvate. SKCO1 cells were maintained in MEM supplemented with 1mM Sodium Pyruvate. Capan-1 cells were maintained in Iscove’s MDM. Calu-1 cells were maintained in McCoy’s 5a medium. MCF10A cells were grown in ATCC’s Mammary Epithelial Cell Basal Medium (ATCC PCS-600-030) supplemented with the Mammary Epithelial Cell Growth Kit (ATCC PCS-600-040) All cells were kept at 37°C in a 5% CO_2_ incubator.

### Compounds

MEK1/2 degraders MS432, MS910 and MS934 have been previously described (Hu et al., 2020, Wei et al., 2019). All other compounds are listed in Data File S2.

### Plasmids

pLKO-Tet-puro-hRAF1-shRNA-1 was a gift from Ayaz Najafov (Addgene plasmid # 185371; http://n2t.net/addgene:185371; RRID:Addgene_185371), MEK1-GFP was a gift from Rony Seger (Addgene plasmid # 14746; http://n2t.net/addgene:14746; RRID:Addgene_14746) pEF-Myc-RAF1 was a gift from Walter Kolch, Myc-Mek1 was a gift from Dustin Maly (Addgene plasmid # 40774; http://n2t.net/addgene:40774; RRID:Addgene_40774) Myc-Mek2 was a gift from Dustin Maly (Addgene plasmid # 40776; http://n2t.net/addgene:40776; RRID:Addgene_40776), mCherry-RAF1 (EX-I5873-Lv111-RAF1), Flag-RAF1 (EX-T0483-M11) Flag-BRAF (EX-Z0707-M11), and Flag-ARAF (EX-B0257-M11) were purchased from Genecopoeia. Mutations were made using the QuikChange XL Site-Directed Mutagenesis Kit (Agilent) according to manufacturer’s protocol. A list of all mutagenic primers is in Data File S2.

### Western Blotting

Samples were harvested in lysis buffer (50 mM HEPES (pH 7.5), 0.5% Triton X-100, 150 mM NaCl, 1 mM EDTA, 1 mM EGTA, 10 mM sodium fluoride, 2.5 mM sodium orthovanadate, 1X protease inhibitor cocktail (Roche), and 1x each of phosphatase inhibitor cocktails 2 and 3 (Sigma)). Particulate was removed by centrifugation of lysates at 21,000 rpm for 15 minutes at 4°C. Lysates were subjected to SDS-PAGE chromatography and transferred to PVDF membranes before western blotting with primary antibodies (**Table S2)**. Secondary HRP-anti-rabbit and HRP-anti-mouse were obtained from ThermoFisher Scientific. SuperSignal West Pico and Femto Chemiluminescent Substrates (Thermo) were used to visualize blots. Western blot images were quantified using the Analyze>Gels function in Image J open-source software (National Institutes of Health). Optical density values for total protein levels were normalized by GAPDH. DC50 (the concentration where 50% of protein has been degraded) plots and values were calculated using PRISM software. Samples were run in biological triplicates (N=3) and Student’s t-tests were performed for statistical analyses and *P* values ≤ 0.05 were considered significant.

### Immunoprecipitations

For Myc- and Flag-tagged protein IPs, cells were transfected 48 hours prior to harvest using Lipofectamine 3000 according to manufacturer’s instructions. Cells were harvested in a buffer containing 25 mM TrisCl (pH 7.5), 1% NP-40, 150 mM NaCl, 1 mM EDTA, 5% glycerol, 1X protease inhibitor cocktail (Roche), and 1x each of phosphatase inhibitor cocktails 2 and 3 (Sigma). Lysates were incubated for 1 hour using over-end rotation, then centrifuged at 21,000 rpm for 15 minutes at 4°C. 1 mg protein was added to 10 μl anti-c-myc or anti-DYKDDDDK magnetic beads (Pierce) and incubated for 2 hours using over-end rotation. Beads were washed 4x with lysis buffer, then boiled in 2x LSB at 100 °C for 5 minutes. For endogenous IPs, cells were harvested as above, then 1 mg protein was precleared using normal mouse IgG agarose conjugate (Santa Cruz) for 2-4 hours at 4°C. Following pre-clearing, lysates were added to 20 μl ARAF, BRAF, RAF1 or MEK1 agarose conjugate (Santa Cruz, Table S2) or normal mouse IgG agarose conjugate and incubated overnight. Beads were washed and proteins eluted as above. TUBE1-IPs were performed as previously described (Alabi et al., 2021). Briefly, cells were harvested in buffer containing 50 mM Tris-HCl, pH 7.4, 0.15 M NaCl, 1 mM EDTA, 1% NP40, 10% glycerol, 5 mM 1,10-phenanthroline monohydrate, 10 mM N-ethylmaleimide, 20 µM PR-619, 1X protease inhibitor cocktail (Roche) and 1x each of phosphatase inhibitor cocktails 2 and 3 (Sigma). 1mg protein was added to 20μl of TUBE1 agarose (LifeSensors) and incubated for 2 hours at 4 °C. Beads were washed 4 times in Tween-TBS and boiled in 2x LSB at 100 °C for 5 minutes. **qRT-PCR**

GeneJET RNA purification kit (Thermo Scientific) was used to isolate RNA from cells according to manufacturer’s instructions. qRT-PCR on diluted cDNA was performed with inventoried TaqMan Gene Expression Assays on the Applied Biosystems QuantStudio™ 6 Real-Time PCR System. The TaqMan Gene Expression Assay probes (ThermoFisher Scientific) used to assess changes in gene expression include *RAF1* (Assay ID: Hs00234119_m1), and ACTB (control) (Cat # 4326315E). Samples were run in biological triplicates (N=3) and Student’s t-tests were performed for statistical analyses and *P* values ≤ 0.05 were considered significant.

### RNAi-mediated Knockdown

siRNA transfections were performed using 25 nM siRNA duplex and the reverse transfection protocol. Cells were added to plates with media containing the siRNA (**Table S2**) and transfection reagent (Lipofectamine RNAiMax) according to the manufacturer’s instructions. Cells were collected 72 h post-transfection. Medium was changed and drug was added 24h prior to collection. Two to three independent experiments were performed with each cell line and siRNA. Samples were run in biological triplicates (N=3) and Student’s t-tests were performed for statistical analyses and *P* values ≤ 0.05 were considered significant.

### Growth and Caspase Assays

For short-term growth assays, 1000-5000 cells were plated per well in 96-well plates and allowed to adhere and equilibrate overnight. Drug was added the following morning and after 120 h of drug treatment, cell viability was assessed using the CellTiter-Glo Luminescent cell viability assay according to manufacturer (Promega). Samples were run in biological triplicates (N=3) and Student’s t-tests were performed for statistical analyses and p values ≤ 0.05 were considered significant. Bliss synergy scores were determined using SynergyFinder (Zheng et al., 2022). Caspase 3/7 activity was determined using the Caspase-Glo 3/7 Assay (Promega) according to manufacturer’s protocol. Cells were treated for 24 hours prior to the assay and luminescence was read 30 minutes following the addition of the reagent.

### Soft Agar Assay

Three ml bottom layer of 0.6% noble agar in RPMI was added per well of a 6 well tissue culture plate. 0.6% noble agar was made by melting a stock solution of 3% noble agar and diluting it 1:5 (600 μl in 2.4 ml media per well) in the respective media and adding it quickly to the plate before it solidifies. Once the bottom layer was solid and cooled (about 20 minutes room temperature), the top layer was added. The top layer was made by adding 300 μl in 1.7 ml media (2 ml total) and storing it in a 15 ml tube in a 42 °C water bath until the cells were ready to be added. Meanwhile, the cells to be added (15,000 HCT-116 or 25,000 A549 cells/well) were placed in a final volume of 1 ml media at 37 °C. The cells were quickly mixed with the 2 ml agar mix (final volume of 3 ml 0.3% agar per well) and added on top of the bottom layer, which was allowed to solidify for at least 30 minutes at room temperature. Once the top layer solidified, 2 ml growth media plus inhibitors or doxycycline was added on top of the top layer. The cells were returned to the incubator and the media was changed every 3-4 days. Cells were allowed to grow for two weeks, then imaged using an EVOS digital inverted microscope at 4x magnification. Following imaging, cells were stained with a 0.1% Crystal Violet solution in 10% ethanol and destained with multiple deionized water washes. Colony number and size were counted using the Analyze Particles function in ImageJ.

### Fluorescence Microscopy

PANC-1 cells were transfected with either mCherry-RAF1 or MEK1-GFP plasmids, then treated with 1 μM MS934 for 24 hours. Images were taken using an EVOS digital inverted microscope at 20x magnification. Images presented in Figure 1 are representative of biological duplicates analyzed.

### Mass Spectrometry and Data Analysis

#### Single Run Total Proteomics and Nano LC MS/MS

DMSO or drug treated cells were lysed in a buffer containing 50 mM HEPES pH 8.0 + 4% SDS, and 100 μg of protein was digested using LysC for 3 hours at room temperature and then trypsin digested overnight at 37°C. Digested peptides were isolated using C-18 stage tips, then dried and cleaned with ethyl acetate. Three μg of proteolytic peptides were resuspended in 0.1% formic acid and separated with a Thermo Scientific RSLCnano Ultimate 3000 LC on a Thermo Scientific Easy-Spray C-18 PepMap 75 µm x 50 cm C-18 2 μm column. A 305 min gradient of 2-20% (180 min) 20%-28% (45 min) 28%-48% (20 min) acetonitrile with 0.1% formic acid was run at 300 nL/min at 50C. Eluted peptides were analyzed by a Thermo Scientific Orbitrap Exploris 480 mass spectrometer utilizing a top 30 methodology in which the 30 most intense peptide precursor ions were subjected to fragmentation. The normalized AGC target for MS1 was set to 300% with a max injection time of 25 ms, the normalized AGC target for MS2 ions was set to 100% with a max injection time of 22 ms, and the dynamic exclusion was set to 90 s.

#### Proteomics Data Processing

Raw data analysis of LFQ experiments was performed using MaxQuant software 2.6.0.1 and searched using the built in Andromeda search engine against the Swiss-Prot human protein database (downloaded on April 24, 2019, 20402 entries). The search was set up for full tryptic peptides with a maximum of two missed cleavage sites. All settings were default and searched using acetylation of protein N-terminus and oxidized methionine as variable modifications. Carbamidomethylation of cysteine was set as a fixed modification. LFQ quantitation was performed using MaxQuant with the following parameters: LFQ minimum ratio count: Global parameters for protein quantitation were as follows: label minimum ratio count: 1, peptides used for quantitation: unique, only use modified proteins selected and with normalized average ratio estimation selected. Samples overexpressing Flag-RAF1 run under the same LC/MS settings were included in the search to assist in detecting lower abundance endogenous RAF1 peptides in the control or drug-treated cells. Match between runs was employed for LFQ quantitation and the significance threshold of the ion score was calculated based on a false discovery rate of < 1%. MaxQuant normalized LFQ values were imported into Perseus software (2.0.11) and filtered in the following manner: Proteins identified by site only were removed, reverse, or potential contaminant were removed then filtered for proteins identified by >1 unique peptide. Protein LFQ values were log2 transformed, filtered for a minimum percent in runs (100%), annotated, and subjected to a Student’s *t*-test comparing treatment *vs.* DMSO. Samples were run in triplicate (N=3). Parameters for the Student’s *t*-test were the following: S0=0.1, two-sided using Permutation-based FDR <0.05. Volcano plots depicting differences in protein abundance were generated using R studio software and Prism graphics. The mass spectrometry proteomics data have been deposited to the ProteomeXchange Consortium via the PRIDE (Vizcaíno et al., 2013) partner repository with the dataset identifier PXD061273 and 10.6019/PXD061273. Reviewer access details: Project accession: PXD061273 Token: qiUcQQybJRqR.

## SUPPLEMENTARY INFORMATION

Figures S1– S7

Data File S1- Raw Label-Free Quantitation (LFQ) values for PANC-1 cells treated with 1 μM MS934 (4 and 24 h) or 1 μM PD0325901 (24 h).

Data File S2- List of reagents.

## Author contributions

J.S.D. contributed experimental design and wrote the manuscript. A.M.K. contributed to manuscript writing and performed Mass spectrometry, immunoblotting, immunoprecipitations, microscopy, qRT-PCR, mutagenesis, stable cell line generation, RNAi knockdown, Cell Titer Glo assays, Caspase Glo assays, 3D soft agar assays. A.M.K. analyzed proteomics datasets and submitted to PRIDE. J.S.W. contributed to manuscript writing and performed immunoblotting and RNAi knockdown. C.H.M. performed immunoblots and immunoprecipitations. G.A.D. contributed to manuscript writing and performed immunoblotting. I.K. and B.F. performed immunoblots. J.J., X.H., and J.H., provided MEK1/2 PROTACs MS432, MS910, and MS934.

## Supplemental Figures

**Figure S1.**
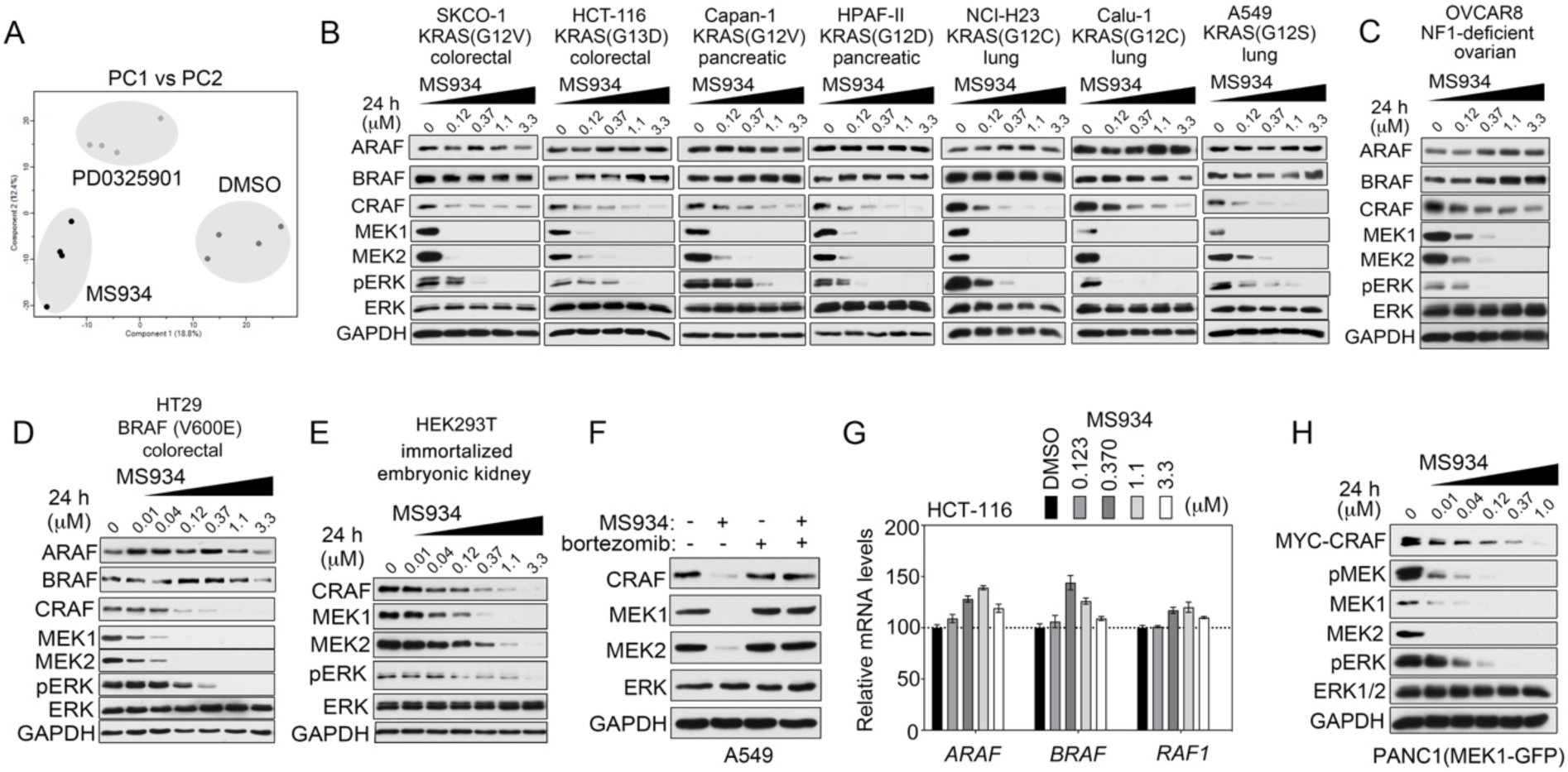
MEK1/2 degradation causes downregulation of CRAF protein levels. (A) Principal component analysis (PCA) of global proteome responses in PANC-1 cells treated with DMSO, or 1 μM PD0325901 or MS934 for 24 h as determined by single-run proteome analysis. (B-E) Immunoblot analysis of MAPK signaling in KRAS mutant cell lines (B) NF1-deficient (C) BRAF mutant (D) or non-tumorigenic cell line (E) treated with escalating doses of MS934 for 24 h. (F) Immunoblotting for CRAF, MEK1/2, or ERK1/2 in A549 treated with DMSO, 0.1 μM MS934, 0.1 μM bortezomib, or the combination for 24 hours. Cells were treated with bortezomib 2 hours prior to administering MS934. (G) *ARAF*, *BRAF*, *RAF1* mRNA levels in HCT-116 cells following treatment with escalating doses of MS934 for 24 hours, as determined by qRT-PCR. Data are means ± SEM of three independent experiments. (H) Immunoblot analysis of Myc-CRAF levels in PANC-1 cells stably expressing MEK1-GFP treated with escalating doses of MS934 for 24 h.

**Figure S2.**
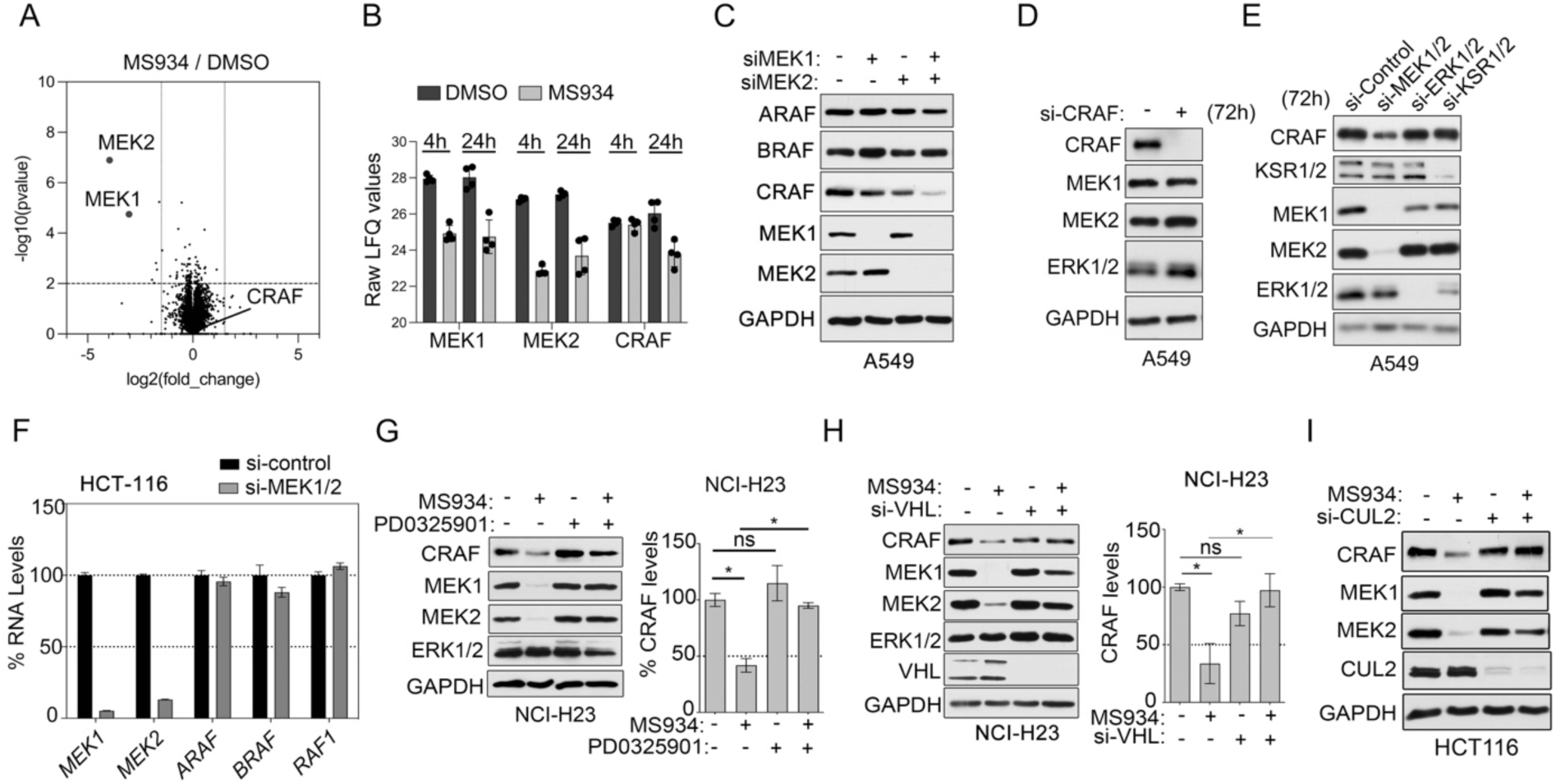
MEK1/2 degradation and knockdown reduce CRAF protein expression. (A) Volcano plot of proteins increased or reduced in abundance in PANC-1 cells treated with MS934 for 4 hours relative to DMSO-treated cells. Differences in protein log2 LFQ intensities among treated or control cells were determined by paired *t-*test permutation-based adjusted *P* values at FDR of <0.05 using Perseus software. (B) Differences in raw LFQ values for MEK1, MEK2, and CRAF in response to DMSO or MS934 treatment for 4 or 24 hours. (C) Immunoblot analysis of RAF and MEK1/2 protein levels in A549 cells treated with control siRNAs or MEK1, MEK2, or MEK1 and MEK2 siRNAs and cultured for 72 hours. (D-E) Immunoblot analysis of CRAF levels in A549 cells treated with non-targeting siRNAs or siRNAs targeting CRAF (A) or MEK1/2, ERK1/2, or KSR1/2 for 72 h. (F) *MEK1/2*, *ARAF*, *BRAF*, *RAF1* mRNA levels in HCT-116 cells following treatment with non- targeting siRNAs or siRNAs targeting MEK1 and MEK2 for 72 hours, as determined by qRT-PCR. Data are means ± SD of three independent experiments. (G) Immunoblotting for CRAF, MEK1/2, or ERK1/2 in NCI-H23 cells treated with DMSO, 1 μM PD0325901, 0.1 μM MS934, or the combination for 24 hours. Cells were treated with PD0325901 2 hours prior to adding MS934. Blots are representative, and densitometric analysis of CRAF levels are means ± SEM from three blots, each normalized to the loading control, GAPDH. (H) Immunoblot analysis for CRAF, MEK1/2, or ERK1/2 in NCI-H23 cells transfected with non- targeting siRNA or siRNAs targeting VHL and treated with DMSO or 0.1 μM MS934 for 24 hours. Blots are representative, and densitometric analysis of CRAF protein levels are means ± SD from three blots, each normalized to the loading control, GAPDH. (I) Immunoblot analysis for CRAF, or MEK1/2 in HCT-116 cells transfected with non-targeting siRNA or siRNAs targeting CUL2 and treated with DMSO or 0.1 μM MS934 for 24 hours.

**Figure S3.**
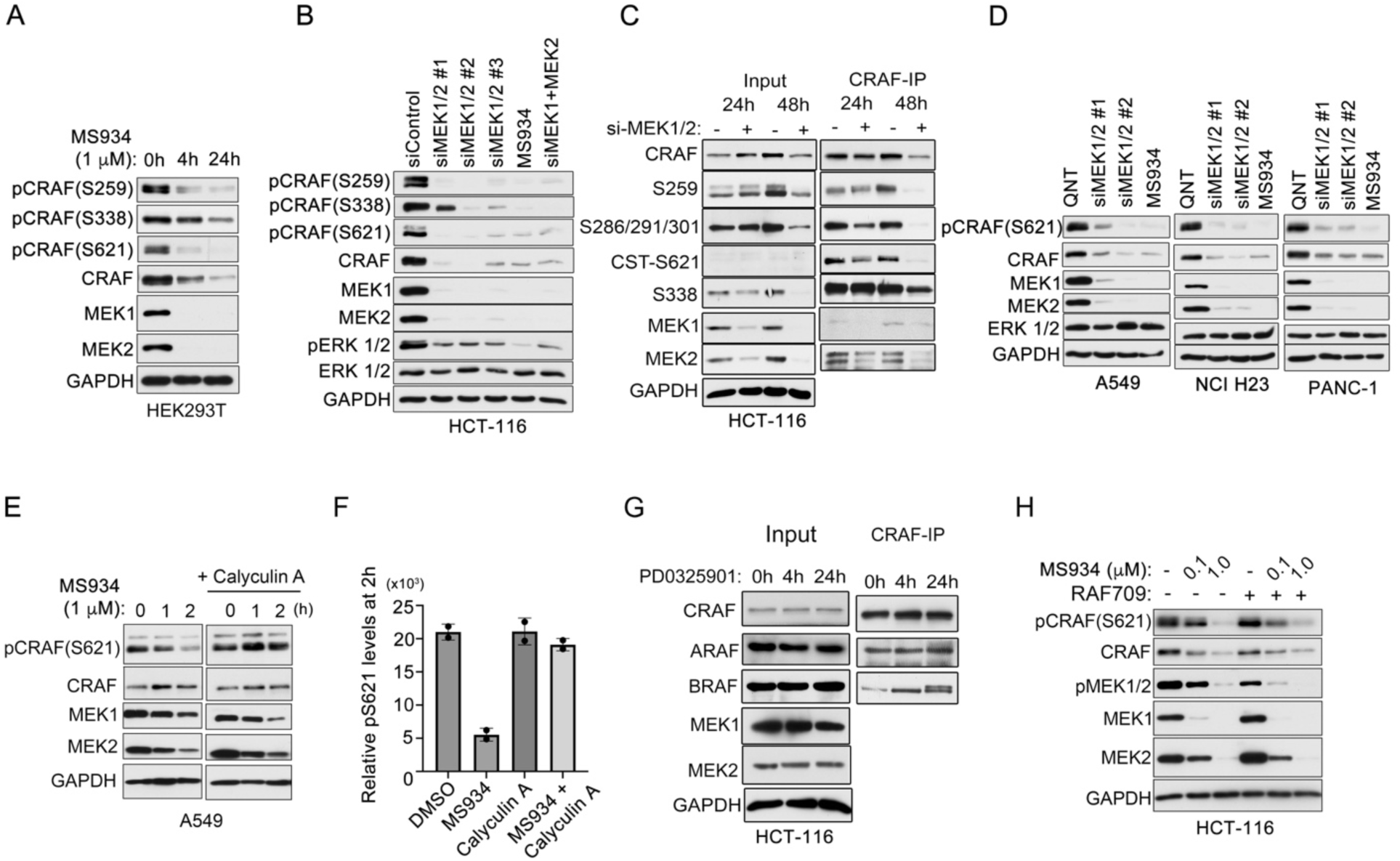
MEK1/2 is essential for preserving the phosphorylation of CRAF preventing its degradation. (A) Immunoblot analysis of CRAF phosphorylation in HEK293T cells treated with 1 μM MS934 for 4 or 24 hours. (B) Immunoblotting for CRAF phosphorylation in HCT-116 cells treated with non-targeting siRNAs, siRNAs targeting a shared sequence amongst MEK1/2, siRNAs targeting MEK1 and MEK2 for 72 h or 1 μM MS934 for 24 h. (C) Immunoprecipitation (IP) of endogenous CRAF from HCT-116 cells treated with non-targeting siRNAs or siRNAs targeting a shared sequence amongst MEK1/2 (siMEK1/2#1) for 24 or 48 h. The CRAF IPs were blotted with the indicated antibodies. Input samples were used as controls. The Cell Signaling Technology (CST) CRAF p-S621 was used. (D) Immunoblot analysis of CRAF phosphorylation in A549, NCI-H23, or PANC-1 cells treated with non-targeting siRNAs or siRNAs targeting a shared sequence amongst MEK1/2 (siMEK1/2#1 or siMEK1/2#2) for 72 h. (E-F) Immunoblotting of CRAF phosphorylation at S621 levels in A549 cells treated with 1 μM MS934 alone or in combination with 2.5 nM Calyculin A for 0, 1, or 2 h. Cells pre-treated with Calyculin A for 30 minutes followed by MS934 treatment for indicated times. Blots are representative, and densitometric analysis of CRAF p-S621 (F) are means ± SEM of 2 independent blots, each normalized to total CRAF protein and the loading control, GAPDH. (G) Immunoprecipitation (IP) of endogenous CRAF from HCT-116 cells treated with 1 μM PD0325901 for 24 h. The CRAF IPs were blotted with the indicated antibodies. Input samples were used as controls. The GAPDH, CRAF, and MEK1/2 input samples are also shown in Fig. 3H, as they were analyzed at the same time. (H) Immunoblot analysis of CRAF total and p-S621, as well as MEK1/2 total and phosphorylated levels in HCT-116 cells treated with 0, 0.1 and 1 μM MS934 alone or combined with 1μM RAF709 for 24 hours.

**Figure S4.**
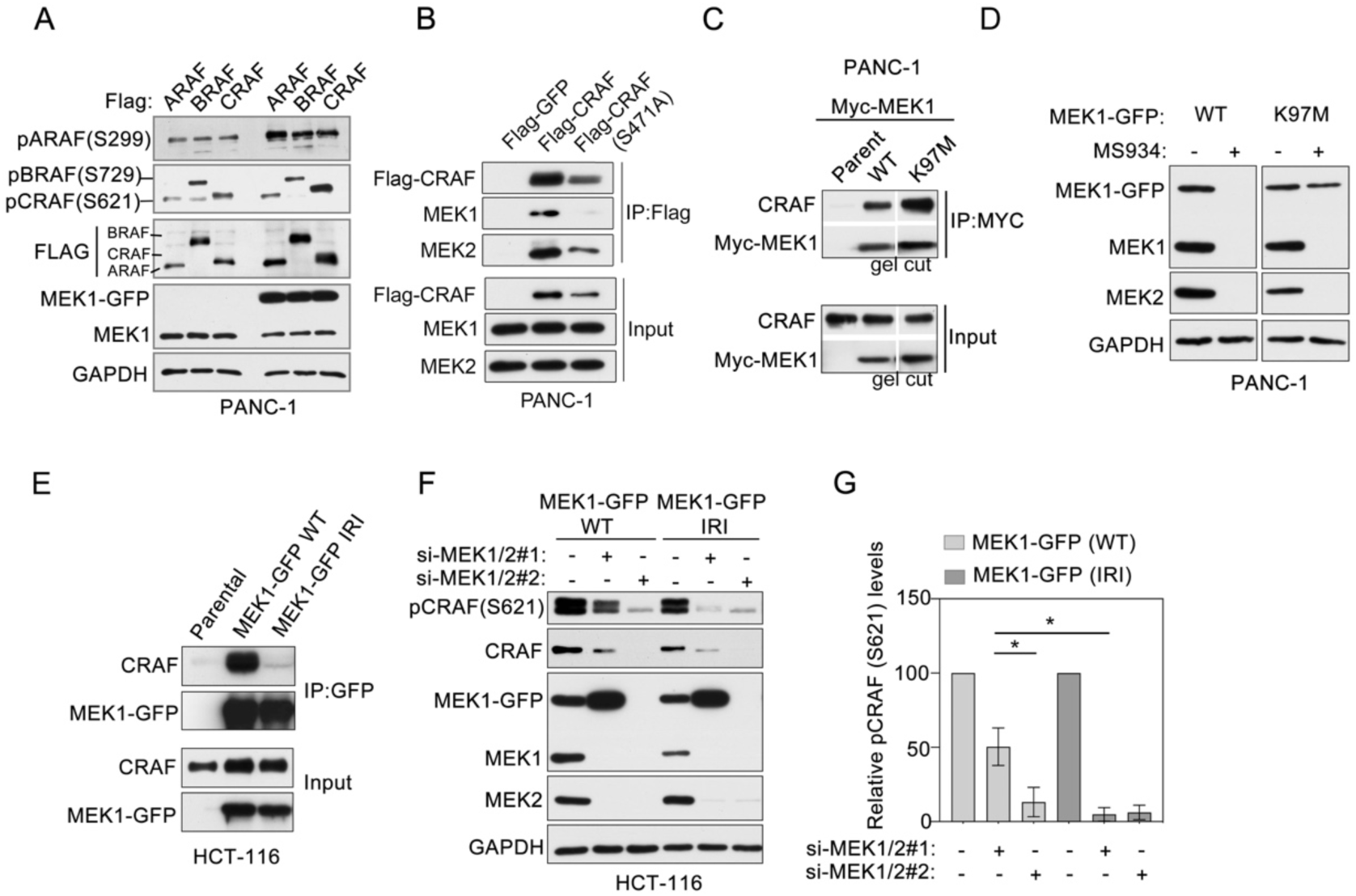
MEK1/2-binding to CRAF and not MEK1/2 kinase activity is required for promoting CRAF phosphorylation at S621 and protein stability. (A) Immunoblot analysis of RAF phosphorylation in PANC-1 or PANC-1 cells stably expressing MEK1-GFP. Flag-ARAF, Flag-BRAF, or Flag-CRAF (all 2.5μg) were transfected into PANC-1 cells. (B) Immunoprecipitation (IP) of Flag-tagged CRAF constructs from PANC-1 cells stably expressing MEK1-GFP. The Flag-CRAF IPs were blotted with the indicated antibodies. IP of Flag-GFP and input samples were used as controls. (C) Immunoprecipitation (IP) of Myc-MEK1 constructs from PANC-1 cells. The Myc-MEK1 IPs were blotted with the indicated antibodies. Input samples were used as controls. (D) Immunoblot analysis of MEK1-GFP levels in PANC-1 cells expressing MEK1-GFP or MEK1- GFP(K97M) treated with 1 μM MS9343 for 24 hours. (E) Immunoprecipitation (IP) of MEK1-GFP constructs from PANC-1 cells. The GFP IPs were blotted with the indicated antibodies. Input samples were used as controls. (F-G) Immunoblot analysis of endogenous CRAF total and p-S621 levels in HCT-116 cells expressing si-MEK1/2#1 resistant MEK1-GFP(WT) o MEK1-GFP(IRI) treated with control, si- MEK1/2#1 or MEK1/2#2 siRNAs for 72 hours. Blots are representative, and densitometric analysis of p-S621 (I) are means ± SEM from three blots, each normalized to the loading control, GAPDH.

**Figure S5.**
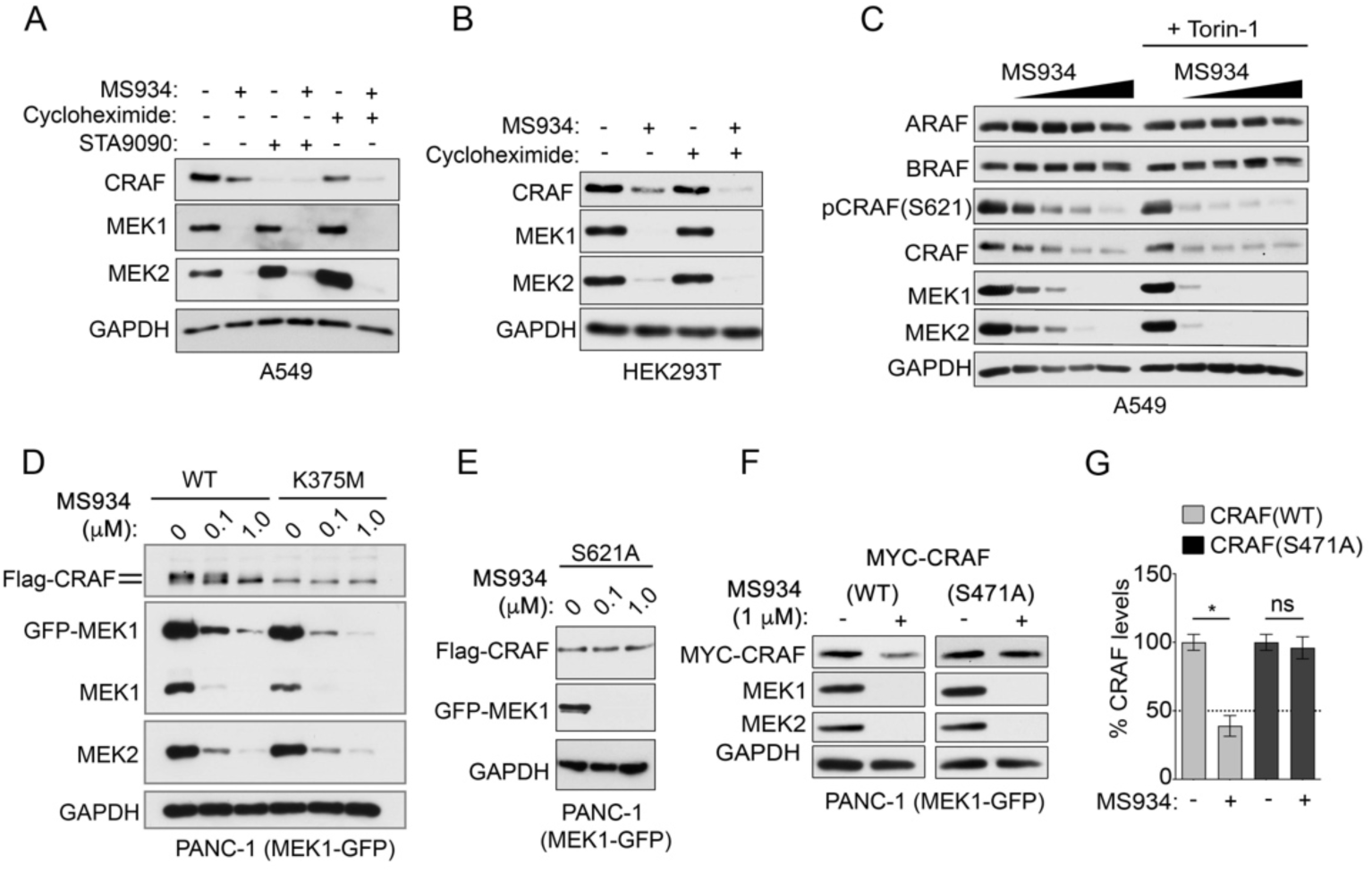
MEK1/2 is required for the stabilization of mature but not immature newly synthesized CRAF which has anti-apoptosis functions. (A) Immunoblotting for CRAF levels in A549 cells treated with DMSO, 1 μM MS934, 100 μg/ml cycloheximide, 0.05 μM STA9090, the combination of MS934 and cycloheximide or STA9090 for 24 h. (B) Immunoblot analysis of CRAF levels in HEK293T cells with DMSO, 1 μM MS934, 100 μg/ml cycloheximide, or the combination of MS934 and cycloheximide for 24 h. (C) Immunoblot analysis for p-S621 and CRAF levels in A549 cells treated with DMSO or escalating doses of MS934 in the presence or absence of 1 μM Torin-1 for 24 hours. (D-E) Immunoblot analysis of Flag-CRAF(WT) or Flag-CRAF(K375M) (D) or Flag-CRAF(S621A) (E) in PANC-1 cells stably expressing MEK1-GFP treated with 0, 0.1 or 1 μM MS934 for 24 h. (F-G) Immunoblot analysis for Myc-CRAF or MEK1/2 in PANC-1 cells stably expressing MEK1- GFP transfected with Myc-CRAF(WT) or Myc-CRAF(S471A) treated with 3.3 μM MS934 for 24 h. Blots are representative, and densitometric analysis of Myc-CRAF levels (G) are means ± SEM from three blots, each normalized to the loading control, GAPDH.

**Figure S6.**
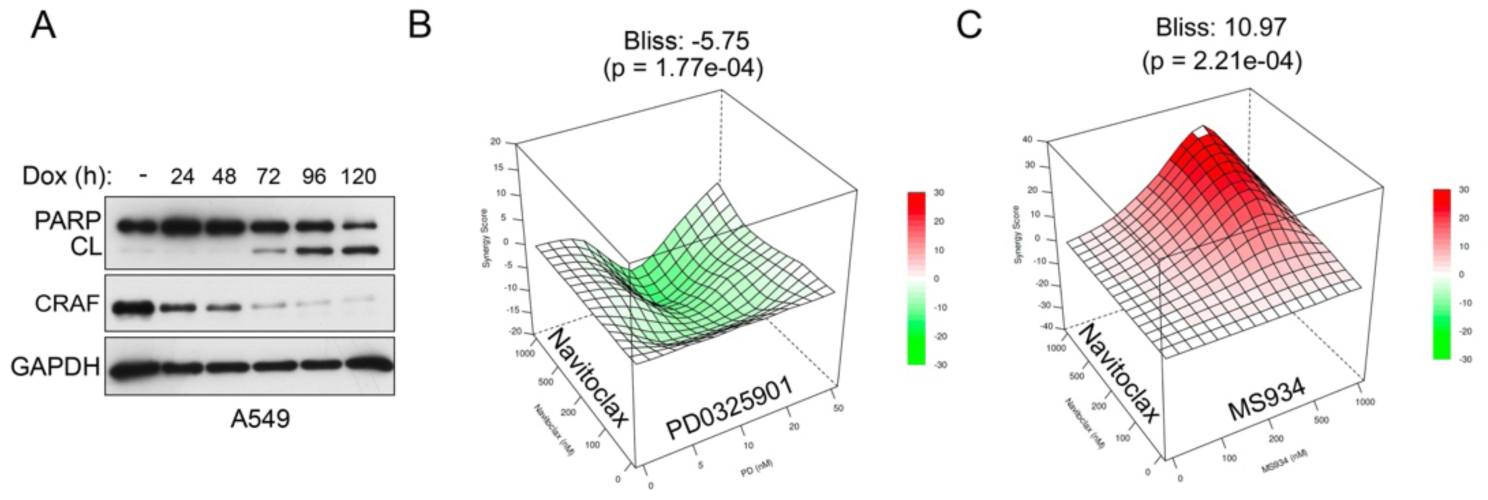
In the absence of MEK1/2 immature CRAF has anti-apoptosis functions. (A) Immunoblot analysis of cleaved PARP levels in A549 cells following short hairpin RNA (shRNA)-mediated depletion of CRAF (shCRAF) or uninduced (1.0 μg/ml doxycycline for 96 h). (B-C) Bliss synergy score based on cell viability by CellTiter-Glo assay of HCT-116 cells treated with escalating doses of Navitoclax, PD0325901 (M), MS934 (N) or the combination for 5 days. Data were analyzed based on % of DMSO control, as means ± SEM of three independent assays.

**Figure S7.**
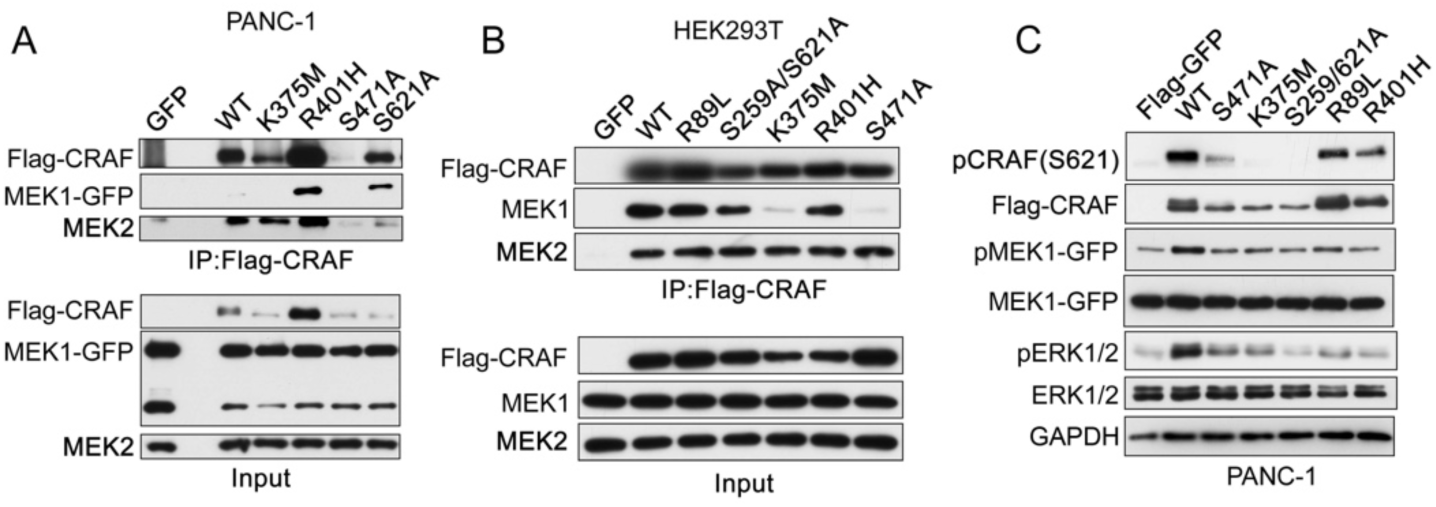
MEK1/2 bind to CRAF prior to RAS engagement forming a primed complex. (A) Immunoprecipitation (IP) of Flag-tagged CRAF constructs from PANC-1 cells stably expressing MEK1-GFP. The Flag-CRAF IPs were blotted with the indicated antibodies. IP of Flag-GFP and input samples were used as controls. (B) Immunoprecipitation (IP) of Flag-tagged CRAF constructs from HEK293T cells stably expressing MEK1-GFP. The Flag-CRAF IPs were blotted with the indicated antibodies. IP of Flag-GFP and input samples were used as controls. (C) Immunoblot analysis of pERK1/2 levels in PANC-1 (MEK1-GFP) cells transfected with Flag- tagged CRAF constructs.

## References

Advani, S. J., Camargo, M. F., Seguin, L., Mielgo, A., Anand, S., Hicks, A. M., Aguilera, J., Franovic, A., Weis, S. M. & Cheresh, D. A. 2015. Kinase-independent role for CRAF-driving tumour radioresistance via CHK2. Nat Commun, 6, 8154.

Alabi, S., Jaime-Figueroa, S., Yao, Z., Gao, Y., Hines, J., Samarasinghe, K. T. G., Vogt, L., Rosen, N. & Crews, C. M. 2021. Mutant-selective degradation by BRAF-targeting PROTACs. Nature Communications, 12, 920.

Bahar, M. E., Kim, H. J. & Kim, D. R. 2023. Targeting the Ras/Raf/Mapk pathway for cancer therapy: from mechanism to clinical studies. Signal Transduct Target Ther, 8, 455.

Boned Del Rio, I., Young, L. C., Sari, S., Jones, G. G., Ringham-Terry, B., Hartig, N., Rejnowicz, E., Lei, W., Bhamra, A., Surinova, S. & Rodriguez-Viciana, P. 2019. SHOC2 complex-driven Raf dimerization selectively contributes to ERK pathway dynamics. Proc Natl Acad Sci U S A, 116, 13330–13339.

Bromberg-White, J. L., Andersen, N. J. & Duesbery, N. S. 2012. Mek genomics in development and disease. Brief Funct Genomics, 11, 300–10.

Cargnello, M. & Roux, P. P. 2012. Activation and Function of the MAPKs and Their Substrates, the Mapk-Activated Protein Kinases. Microbiology and Molecular Biology Reviews, 76, 496–496.

Catalanotti, F., Reyes, G., Jesenberger, V., Galabova-Kovacs, G., De Matos Simoes, R., Carugo, O. & Baccarini, M. 2009. A Mek1-Mek2 heterodimer determines the strength and duration of the Erk signal. Nat Struct Mol Biol, 16, 294–303.

Chen, J., Fujii, K., Zhang, L., Roberts, T. & Fu, H. 2001. Raf-1 promotes cell survival by antagonizing apoptosis signal-regulating kinase 1 through a Mek–Erk independent mechanism. Proceedings of the National Academy of Sciences, 98, 7783–7788.

Dhillon, A. S., Von Kriegsheim, A., Grindlay, J. & Kolch, W. 2007. Phosphatase and feedback regulation of Raf-1 signaling. Cell Cycle, 6, 3–7.

Dougherty, M. K., Müller, J., Ritt, D. A., Zhou, M., Zhou, X. Z., Copeland, T. D., Conrads, T. P., Veenstra, T. D., Lu, K. P. & Morrison, D. K. 2005. Regulation of Raf-1 by Direct Feedback Phosphorylation. Molecular Cell, 17, 215–224.

Ferrier, A. F., Lee, M., Anderson, W. B., Benvenuto, G., Morrison, D. K., Lowy, D. R. & Declue, J. E. 1997. Sequential modification of serines 621 and 624 in the Raf-1 carboxyl terminus produces alterations in its electrophoretic mobility. J Biol Chem, 272, 2136–42.

Fujioka, A., Terai, K., Itoh, R. E., Aoki, K., Nakamura, T., Kuroda, S., Nishida, E. & Matsuda, M. 2006. Dynamics of the Ras/Erk MAPK cascade as monitored by fluorescent probes. J Biol Chem, 281, 8917–26.

Garcia-Alonso, S., Mesa, P., Ovejero, L. P., Aizpurua, G., Lechuga, C. G., Zarzuela, E., Santiveri, C. M., Sanclemente, M., Munoz, J., Musteanu, M., Campos- Olivas, R., Martinez-Torrecuadrada, J., Barbacid, M. & Montoya, G. 2022. Structure of the RAF1-HSP90-CDC37 complex reveals the basis of RAF1 regulation. Mol Cell, 82, 3438–3452 e8.

Grammatikakis, N., Lin, J. H., Grammatikakis, A., Tsichlis, P. N. & Cochran, B. H. 1999. p50(cdc37) acting in concert with Hsp90 is required for Raf-1 function. Mol Cell Biol, 19, 1661–72.

Grbovic, O. M., Basso, A. D., Sawai, A., Ye, Q., Friedlander, P., Solit, D. & Rosen, N. 2006. V600e B-Raf requires the Hsp90 chaperone for stability and is degraded in response to Hsp90 inhibitors. Proc Natl Acad Sci U S A, 103, 57–62.

Haling, J. R., Sudhamsu, J., Yen, I., Sideris, S., Sandoval, W., Phung, W., Bravo, B. J., Giannetti, A. M., Peck, A., Masselot, A., Morales, T., Smith, D., Brandhuber, B. J., Hymowitz, S. G. & Malek, S. 2014. Structure of the Braf-Mek complex reveals a kinase activity independent role for Braf in MAPK signaling. Cancer Cell, 26, 402–413.

Hatzivassiliou, G., Haling, J. R., Chen, H., Song, K., Price, S., Heald, R., Hewitt, J. F., Zak, M., Peck, A., Orr, C., Merchant, M., Hoeflich, K. P., Chan, J., Luoh, S. M., Anderson, D. J., Ludlam, M. J., Wiesmann, C., Ultsch, M., Friedman, L. S., Malek, S. & Belvin, M. 2013. Mechanism of MEK inhibition determines efficacy in mutant Kras- versus Braf-driven cancers. Nature, 501, 232–6.

Hjerpe, R., Aillet, F., Lopitz-Otsoa, F., Lang, V., England, P. & Rodriguez, M. S. 2009. Efficient protection and isolation of ubiquitylated proteins using tandem ubiquitin-binding entities. Embo reports, 10, 1250–1258-1258.

Hu, J., Wei, J., Yim, H., Wang, L., Xie, L., Jin, M. S., Kabir, M., Qin, L., Chen, X., Liu, J. & Jin, J. 2020. Potent and Selective Mitogen-Activated Protein Kinase Kinase 1/2 (MEK1/2) Heterobifunctional Small-molecule Degraders. J Med Chem, 63, 15883–15905.

Huth, H. W., Albarnaz, J. D., Torres, A. A., Bonjardim, C. A. & Ropert, C. 2016. MEK2 controls the activation of MKK3/MKK6-p38 axis involved in the Mda-Mb-231 breast cancer cell survival: Correlation with cyclin D1 expression. Cellular Signalling, 28, 1283–1291.

Kamal, A., Boehm, M. F. & Burrows, F. J. 2004. Therapeutic and diagnostic implications of Hsp90 activation. Trends Mol Med, 10, 283–90.

King, A. J., Sun, H., Diaz, B., Barnard, D., Miao, W., Bagrodia, S. & Marshall, M. S. 1998. The protein kinase Pak3 positively regulates Raf-1 activity through phosphorylation of serine 338. Nature, 396, 180–3.

Kurimchak, A. M., Herrera-Montávez, C., Montserrat-Sangrà, S., Araiza- Olivera, D., Hu, J., Neumann-Domer, R., Kuruvilla, M., Bellacosa, A., Testa, J. R., Jin, J. & Duncan, J. S. 2022. The drug efflux pump MDR1 promotes intrinsic and acquired resistance to PROTACs in cancer cells. Sci Signal, 15, eabn2707.

Lamba, S., Russo, M., Sun, C., Lazzari, L., Cancelliere, C., Grernrum, W., Lieftink, C., Bernards, R., Di Nicolantonio, F. & Bardelli, A. 2014. Raf Suppression Synergizes with Mek Inhibition in *Kras* Mutant Cancer Cells. Cell Reports, 8, 1475-1483.

Lavoie, H., Gagnon, J. & Therrien, M. 2020. Erk signalling: a master regulator of cell behaviour, life and fate. Nat Rev Mol Cell Biol, 21, 607–632.

Lavoie, H., Sahmi, M., Maisonneuve, P., Marullo, S. A., Thevakumaran, N., Jin, T., Kurinov, I., Sicheri, F. & Therrien, M. 2018. Mek drives Braf activation through allosteric control of Ksr proteins. Nature, 554, 549–553.

Lavoie, H. & Therrien, M. 2015. Regulation of Raf protein kinases in Erk signalling. Nature Reviews Molecular Cell Biology, 16, 281–298.

Leicht, D. T., Balan, V., Zhu, J., Kaplun, A., Bronisz, A., Rana, A. & Tzivion, G. 2013. Mek-1 activates C-Raf through a Ras-independent mechanism. Biochimica et Biophysica Acta (Bba) - Molecular Cell Research, 1833, 976–986.

Lito, P., Saborowski, A., Yue, J., Solomon, M., Joseph, E., Gadal, S., Saborowski, M., Kastenhuber, E., Fellmann, C., Ohara, K., Morikami, K., Miura, T., Lukacs, C., Ishii, N., Lowe, S. & Rosen, N. 2014. Disruption of Craf-mediated Mek activation is required for effective Mek inhibition in Kras mutant tumors. Cancer Cell, 25, 697–710.

Mason, C. S., Springer, C. J., Cooper, R. G., Superti-Furga, G., Marshall, C. J. & Marais, R. 1999. Serine and tyrosine phosphorylations cooperate in Raf-1, but not B-Raf activation. Embo j, 18, 2137–48.

Mccormick, F. 2018. c-Raf in KRas Mutant Cancers: A Moving Target. Cancer cell, 33, 158–159.

Mckay, M. M., Ritt, D. A. & Morrison, D. K. 2009. Signaling dynamics of the KSR1 scaffold complex. Proc Natl Acad Sci U S A, 106, 11022–7.

Mielgo, A., Seguin, L., Huang, M., Camargo, M. F., Anand, S., Franovic, A., Weis, S. M., Advani, S. J., Murphy, E. A. & Cheresh, D. A. 2011. A Mek-independent role for Craf in mitosis and tumor progression. Nat Med, 17, 1641–5.

Mitra, S., Ghosh, B., Gayen, N., Roy, J. & Mandal, A. K. 2016. Bipartite Role of Heat Shock Protein 90 (Hsp90) Keeps Craf Kinase Poised for Activation. J Biol Chem, 291, 24579–24593.

Noble, C., Mercer, K., Hussain, J., Carragher, L., Giblett, S., Hayward, R., Patterson, C., Marais, R. & Pritchard, C. A. 2008. Craf autophosphorylation of serine 621 is required to prevent its proteasome-mediated degradation. Mol Cell, 31, 862–72.

O’neill, E., Rushworth, L., Baccarini, M. & Kolch, W. 2004. Role of the kinase MST2 in suppression of apoptosis by the proto-oncogene product Raf-1. Science, 306, 2267–70.

Ory, S., Zhou, M., Conrads, T. P., Veenstra, T. D. & Morrison, D. K. 2003. Protein phosphatase 2a positively regulates Ras signaling by dephosphorylating KSR1 and Raf-1 on critical 14-3-3 binding sites. Curr Biol, 13, 1356–64.

Pearson, G., Bumeister, R., Henry, D. O., Cobb, M. H. & White, M. A. 2000. Uncoupling Raf1 from MEK1/2 impairs only a subset of cellular responses to Raf activation. J Biol Chem, 275, 37303–6.

Rauch, J., Volinsky, N., Romano, D. & Kolch, W. 2011. The secret life of kinases: functions beyond catalysis. Cell Communication and Signaling, 9, 23.

Reiterer, G., Chen, L., Tassef, R., Varner, J. D., Chen, C. Y. & Yen, A. 2010. Raf associates with phosphorylated nuclear BubR1 during endoreduplication induced by Jak inhibition. Cell Cycle, 9, 3297–304.

Roskoski, R., Jr. 2012. MEK1/2 dual-specificity protein kinases: structure and regulation. Biochem Biophys Res Commun, 417, 5–10.

Sanclemente, M., Nieto, P., Garcia-Alonso, S., Fernandez-Garcia, F., Esteban- Burgos, L., Guerra, C., Drosten, M., Caleiras, E., Martinez- Torrecuadrada, J., Santamaria, D., Musteanu, M. & Barbacid, M. 2021. RAF1 kinase activity is dispensable for Kras/p53 mutant lung tumor progression. Cancer Cell, 39, 294–296.

Santarpia, L., Lippman, S. M. & El-Naggar, A. K. 2012. Targeting the Mapk-Ras-Raf signaling pathway in cancer therapy. Expert Opin Ther Targets, 16, 103–19.

Schulte, T. W., Blagosklonny, M. V., Ingui, C. & Neckers, L. 1995. Disruption of the Raf-1-Hsp90 molecular complex results in destabilization of Raf-1 and loss of Raf-1- Ras association. J Biol Chem, 270, 24585–8.

Shao, W., Mishina, Y. M., Feng, Y., Caponigro, G., Cooke, V. G., Rivera, S., Wang, Y., Shen, F., Korn, J. M., Mathews Griner, L. A., Nishiguchi, G., Rico, A., Tellew, J., Haling, J. R., Aversa, R., Polyakov, V., Zang, R., Hekmat-Nejad, M., Amiri, P., Singh, M., Keen, N., Dillon, M. P., Lees, E., Ramurthy, S., Sellers, W. R. & Stuart, D. D. 2018. Antitumor Properties of RAF709, a Highly Selective and Potent Inhibitor of Raf Kinase Dimers, in Tumors Driven by Mutant Ras or Braf. Cancer Res, 78, 1537–1548.

Silverstein, A. M., Grammatikakis, N., Cochran, B. H., Chinkers, M. & Pratt, W. B. 1998. p50(cdc37) binds directly to the catalytic domain of Raf as well as to a site on hsp90 that is topologically adjacent to the tetratricopeptide repeat binding site. J Biol Chem, 273, 20090–5.

Singh, A. K., Sonawane, P., Kumar, A., Singh, H., Naumovich, V., Pathak, P., Grishina, M., Khalilullah, H., Jaremko, M., Emwas, A. H., Verma, A. & Kumar, P. 2023. Challenges and Opportunities in the Crusade of Braf Inhibitors: From 2002 to 2022. Acs Omega, 8, 27819–27844.

Smith, J. R., Clarke, P. A., De Billy, E. & Workman, P. 2009. Silencing the cochaperone CDC37 destabilizes kinase clients and sensitizes cancer cells to HSP90 inhibitors. Oncogene, 28, 157–69.

Thorson, J. A., Yu, L. W., Hsu, A. L., Shih, N. Y., Graves, P. R., Tanner, J. W., Allen, P. M., Piwnica-Worms, H. & Shaw, A. S. 1998. 14-3-3 proteins are required for maintenance of Raf-1 phosphorylation and kinase activity. Mol Cell Biol, 18, 5229–38.

Venkatanarayan, A., Liang, J., Yen, I., Shanahan, F., Haley, B., Phu, L., Verschueren, E., Hinkle, T. B., Kan, D., Segal, E., Long, J. E., Lima, T., Liau, N. P. D., Sudhamsu, J., Li, J., Klijn, C., Piskol, R., Junttila, M. R., Shaw, A. S., Merchant, M., Chang, M. T., Kirkpatrick, D. S. & Malek, S. 2022. Craf dimerization with Araf regulates Kras-driven tumor growth. Cell Rep, 38, 110351.

Vizcaíno, J. A., Côté, R. G., Csordas, A., Dianes, J. A., Fabregat, A., Foster, J. M., Griss, J., Alpi, E., Birim, M., Contell, J., O’kelly, G., Schoenegger, A., Ovelleiro, D., Pérez-Riverol, Y., Reisinger, F., Ríos, D., Wang, R. & Hermjakob, H. 2013. The PRoteomics IDEntifications (Pride) database and associated tools: status in 2013. Nucleic Acids Res, 41, D1063–9.

Wei, J., Hu, J., Wang, L., Xie, L., Jin, M. S., Chen, X., Liu, J. & Jin, J. 2019. Discovery of a First-in-Class Mitogen-Activated Protein Kinase Kinase 1/2 Degrader. Journal of Medicinal Chemistry, 62, 10897–10911.

Xiang, X., Zang, M., Waelde, C. A., Wen, R. & Luo, Z. 2002. Phosphorylation of 338SSYY341 regulates specific interaction between Raf-1 and MEK1. J Biol Chem, 277, 44996–5003.

Xiong, Y., Zhong, Y., Yim, H., Yang, X., Park, K. S., Xie, L., Poulikakos, P. I., Han, X., Xiong, Y., Chen, X., Liu, J. & Jin, J. 2022. Bridged Proteolysis Targeting Chimera (Protac) Enables Degradation of Undruggable Targets. J Am Chem Soc, 144, 22622–22632.

Yamamoto, K., Ichijo, H. & Korsmeyer, S. J. 1999. Bcl-2 is phosphorylated and inactivated by an ASK1/Jun N-terminal protein kinase pathway normally activated at G(2)/M. Mol Cell Biol, 19, 8469–78.

Yu, A., Nguyen, D. H., Nguyen, T. J. & Wang, Z. 2023. A novel phosphorylation site involved in dissociating Raf kinase from the scaffolding protein 14-3-3 and disrupting Raf dimerization. J Biol Chem, 299, 105188.

Zang, M., Gong, J., Luo, L., Zhou, J., Xiang, X., Huang, W., Huang, Q., Luo, X., Olbrot, M., Peng, Y., Chen, C. & Luo, Z. 2008. Characterization of Ser338 phosphorylation for Raf-1 activation. J Biol Chem, 283, 31429–37.

Zheng, S., Wang, W., Aldahdooh, J., Malyutina, A., Shadbahr, T., Tanoli, Z., Pessia, A. & Tang, J. 2022. SynergyFinder Plus: Toward Better Interpretation and Annotation of Drug Combination Screening Datasets. Genomics, Proteomics & Bioinformatics, 20, 587–596.

Zhu, J., Balan, V., Bronisz, A., Balan, K., Sun, H., Leicht, D. T., Luo, Z., Qin, J., Avruch, J. & Tzivion, G. 2005. Identification of Raf-1 S471 as a novel phosphorylation site critical for Raf-1 and B-Raf kinase activities and for Mek binding. Mol Biol Cell, 16, 4733–44.

